# Integrative analysis of neuroblastoma by single-cell RNA sequencing identifies the NECTIN2-TIGIT axis as a target for immunotherapy

**DOI:** 10.1101/2022.07.15.499859

**Authors:** Judith Wienke, Lindy L. Visser, Waleed M. Kholosy, Kaylee M. Keller, Marta Barisa, Sophie Munnings-Tomes, Elizabeth Carlton, Evon Poon, Ana Rodriguez, Ronald Bernardi, Femke van den Ham, Sander R. van Hooff, Yvette A.H. Matser, Michelle L. Tas, Karin P.S. Langenberg, Philip Lijnzaad, Josephine G.M. Strijker, Alvaro Sanchez-Bernabeu, Annelisa M. Cornel, Frank C.P. Holstege, Juliet Gray, Lieve A.M. Tytgat, Ronald R. de Krijger, Marijn A. Scheijde-Vermeulen, Marc H.W.A. Wijnen, Miranda Dierselhuis, Karin Straathof, Sam Behjati, Wei Wu, Albert J.R. Heck, Jan Koster, Stefan Nierkens, Louis Chesler, John Anderson, Hubert N. Caron, Thanasis Margaritis, Max M. van Noesel, Jan J. Molenaar

## Abstract

Pediatric patients with high-risk neuroblastoma have poor survival rates and urgently need more effective treatment options with less side effects. As novel and improved immunotherapies may fill this need, we dissected the immunoregulatory interactions in neuroblastoma by single-cell RNA-sequencing of 25 tumors (10 pre- and 15 post-chemotherapy, including 5 pairs) to identify strategies for optimizing immunotherapy efficacy. Neuroblastomas were infiltrated by NK, T and B cells, and immunosuppressive myeloid populations. NK cells showed reduced cytotoxicity and T cells had a dysfunctional profile. Interaction analysis revealed a vast immunoregulatory network and identified NECTIN2-TIGIT as a crucial immune checkpoint. Combined blockade of TIGIT and PD-L1 significantly reduced neuroblastoma growth, with complete responses *in vivo*. Moreover, addition of TIGIT blockade to standard relapse treatment in a chemotherapy-resistant *Th*-*ALK*^F1174L^/*MYCN* 129/SvJ syngeneic model significantly improved survival. Concluding, our integrative analysis of neuroblastoma’s vast immunoregulatory network provides novel targets and a rationale for immunotherapeutic combination strategies.

## INTRODUCTION

Immunotherapy has revolutionized cancer treatment in adults and holds great promise for pediatric solid tumors(1). Its potential is exemplified by the increased survival of patients with high-risk neuroblastoma following implementation of anti-GD2 antibody therapy into standard care(2,3). Neuroblastoma, the most common extracranial pediatric solid tumor, accounts for 10% of pediatric cancer-related deaths(4). Patients are stratified into risk groups based on disease presentation and genomic alterations like *MYCN* amplification, which is an important driver of poor prognosis(5). High-risk neuroblastoma patients receive an intense multimodal treatment regimen, consisting of induction chemotherapy, surgical tumor resection, consolidation with high-dose chemotherapy followed by autologous stem cell transplantation, radiotherapy and anti-GD2 immunotherapy. Introduction of anti-GD2 has improved high-risk neuroblastoma event-free survival rates by ∼15%(2,3). Still, overall 5-year survival rates are below 60%, particularly due to the high relapse rate(3). Chemoimmunotherapy with temozolomide, irinotecan and anti-GD2 is likely to be selected as the backbone treatment for relapse/refractory neuroblastoma patients in Europe(6). Due to the intense treatment, the majority of survivors suffer of debilitating (long-term) side effects(7). Taken together, there is an urgent need for novel, effective treatments with less side effects.

The success of anti-GD2 therapy has provided a clear rationale for immunotherapy in neuroblastoma treatment. But, as the survival benefit is still modest, improving immunotherapy efficacy will be crucial. T cells and natural killer (NK) cells are considered essential effectors in the context of immunotherapy, but in many cancers their function is compromised(8,9). Efforts to improve immunotherapy efficacy focus on reinvigorating T and NK cell function, e.g. with immune checkpoint inhibition or by introducing cellular immunotherapies like CAR-T cells(1,10). So far, checkpoint inhibitor monotherapy and CAR-T cells have shown relatively limited efficacy in clinical trials for patients with neuroblastoma and have not yet led to a consistent survival benefit(11–19). This limited success may be due to intra-tumoral factors hampering their efficacy, such as immunosuppressive cells and immune checkpoints(10,20). Efforts to optimize immunotherapies for neuroblastoma are currently hampered by a lack of information on these intra-tumoral factors. The detailed composition and function of T and NK cells in the neuroblastoma tumor-microenvironment is still largely unexplored and only few immunosuppressive factors, cq. immune checkpoint molecules (e.g. PD-L1 and B7-H3), have so far been identified(21). To guide further immunotherapy development – and enhancement – it is essential to gain better insights into neuroblastoma’s immune environment and the immunoregulatory mechanisms at play. Moreover, such insights may fuel an educated approach to design combination immunotherapy strategies overcoming immune resistance(21).

To unravel the immune environment of neuroblastoma and identify novel targets for immunotherapy enhancement, we generated a single-cell transcriptomic atlas of 25 tumors, with samples taken before and after induction chemotherapy. We provide a detailed view of neuroblastoma’s immune landscape pre- and post-chemotherapy, and reveal a vast immunoregulatory interaction network. With subsequent validation experiments we identified the NECTIN2-TIGIT axis as promising target for neuroblastoma immunotherapy.

## RESULTS

### The single-cell landscape of neuroblastoma

To provide an in-depth view of the neuroblastoma’s tumor-microenvironment, samples were analyzed by single-cell RNA sequencing. Fresh material was processed from 25 tumors of 20 patients, 10 taken pre-treatment and 15 after induction chemotherapy, including 5 paired samples. Seventeen patients had high-risk neuroblastoma, 2 medium-risk neuroblastoma, and 1 patient ganglioneuroma, and 6/20 patients had *MYCN* amplified (*MYCN*-A) tumors (Fig. 1a, Supplementary table 1). Single-cell RNA sequencing with deep sequencing Cel-Seq2 protocol yielded 12,232 high-quality cells in total. We identified 21 cell clusters, which were annotated as 5 main cell types, i.e. tumor, immune cells, endothelium, mesenchyme, and Schwann cell precursor cells (Fig. 1b,c and Supplementary Fig. 1a). The three tumor clusters highly expressed *CDK4, CCND1* and *MYCN*, respectively (Fig. 1c). Tumor cluster 1 contained *CDK4* amplified cells, and cluster 3 contained *MYCN*-A cells (Supplementary Table 1). In addition, the three tumor clusters highly and specifically expressed previously described neuroblastoma markers genes, including *CHGA, PHOX2B* and *TH* (Fig. 1d and Supplementary Fig. 1b)(22). Tumor cell identity was confirmed by copy number variation inference, showing typical chromosomal aberrations (e.g. 1p loss, 11q loss and 17q gain; Supplementary Fig. 1c)(22). As reported before(23), HLA class I (*HLA-A, HLA-B, HLA-C*) expression was significantly lower in tumor than non-tumor cells, particularly in tumors with high *MYCN* expression (Supplementary Fig. 1d-g), highlighting neuroblastoma’s low immunogenicity. The cellular composition of tumors differed per patient, but distinct changes were observed upon induction chemotherapy (Fig. 1e and Supplementary Fig. 1h). The tumor content decreased from 47% (range 12-97%) pre-treatment to only 2% (0-17%) post-treatment, while the mesenchymal fraction concordantly increased from 12% (0-49%) to 52% (4-91%) (both p<0.005, Fig. 1f,g). The other main cell fractions, including immune cells, did not change upon treatment (Fig. 1f). Although MYCN-A tumors are associated with low immune cell infiltration(24), their cellular composition did not differ from *MYCN*-non amplified (NA) tumors (Supplementary Fig. 1i).

**Figure 1.**
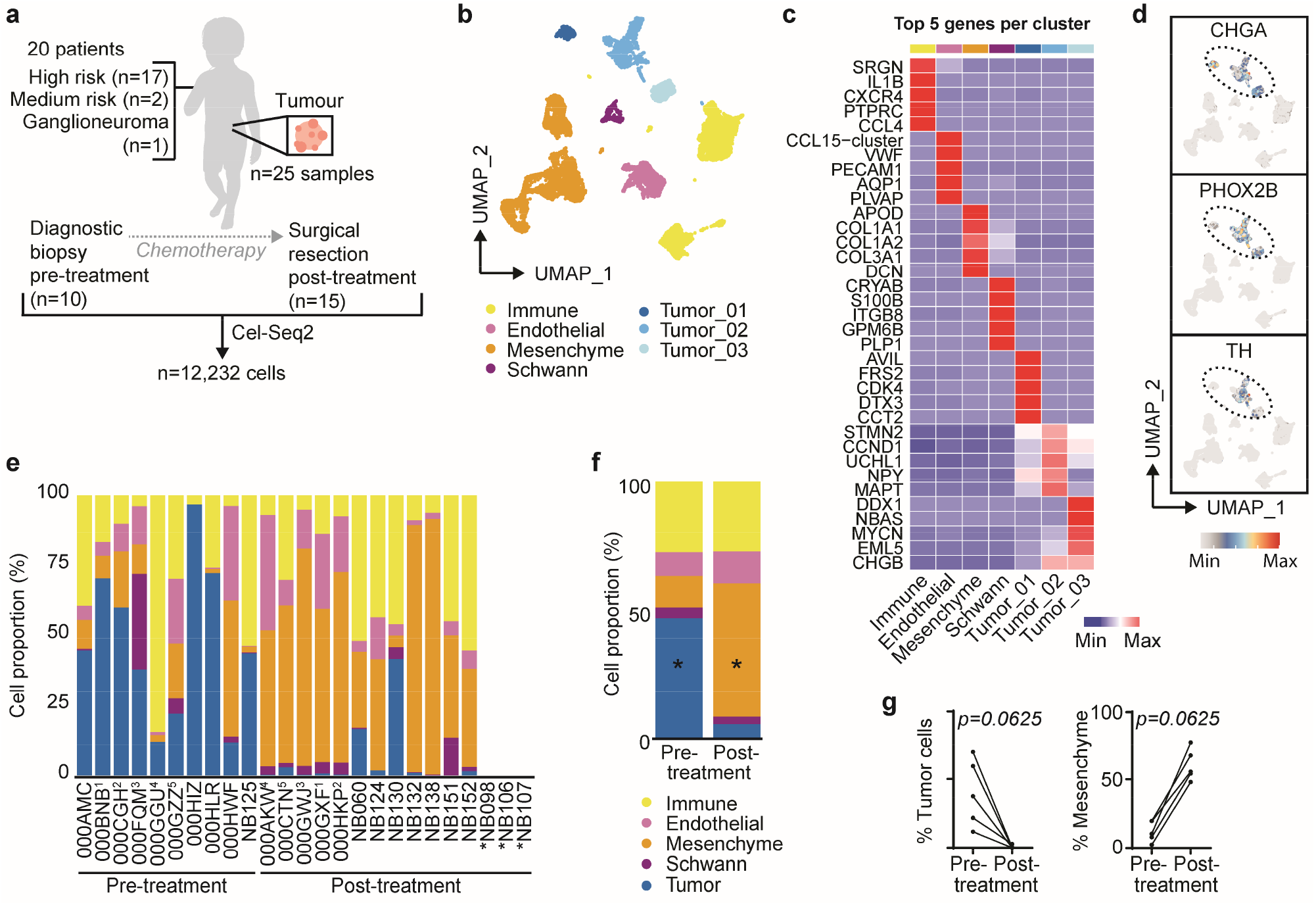
The single-cell landscape of neuroblastoma. (**a**) Schematic overview of included patients and sample collection. (**b**) UMAP of main cell types. (**c**) Heatmap of top 5 differentially expressed genes per main cell type by average log-normalized expression. (**d**) UMAP with expression of typical neuroblastoma genes. (**e**) Proportion of different cell types per sample. *To calculate the proportion, only cells which were sorted with an unbiased sorting strategy were included, which led to exclusion of NB098, NB106 and NB107. ^1,2,3,4,5^: Correspond to paired samples before and after treatment. (**f**) Average cellular composition of samples before and after induction chemotherapy. Mixed-effects analysis with Sidak’s multiple comparisons test. *\*P<0*.*005*. (**g**) Cell proportion of mesenchymal and tumor cells in five paired pre-treatment and post-treatment samples.

### Neuroblastoma is populated by immunosuppressive macrophages

To unravel the composition and function of neuroblastoma-infiltrating immune cells we performed an in-depth analysis of the *PTPRC*^+^ (CD45^+^) immune clusters, which identified, among a total of 3,602 immune cells, myeloid and lymphoid cells (Fig. 2a and Supplementary Fig. 2a,b). The myeloid cells, which are considered key regulators of anti-tumor immunity due to their role in antigen presentation and T cell activation, but also immunosuppression(25), consisted of mast cells, plasmacytoid dendritic cells (pDC), conventional DC (cDC), *S100A8/A9*^hi^ undifferentiated monocytes (Mo), and four differentiated macrophage populations (*IL10*^hi^, *APO*^hi^, *CCL2*^hi^ and *MAF*^hi^ macrophages (Mϕ; Fig. 2b). cDC, and *IL10*^hi^, *APO*^hi^ and *MAF*^hi^ Mϕ had the highest expression of HLA and costimulatory molecules (p<0.0001 vs *S100*^hi^ Mo), rendering them the most potent interaction partners for T cell (co)-stimulation (Fig. 2c and Supplementary Fig. 2c). Compared to the undifferentiated *S100*^hi^ Mo, all macrophage populations displayed an M2-like signature associated with immunosuppressive and pro-tumorigenic properties (p<0.0001) (Fig. 2d)(26,27). The *IL10*^hi^ Mϕ population had a mixed profile, expressing M1-like (pro-inflammatory) features next to M2-like characteristics (p<0.0001; Fig. 2d). The immunoregulatory nature of the different myeloid populations was further substantiated by their expression of soluble factors with acknowledged immunosuppressive roles in tumors, such as *IL10, LGALS3*, and *MMP9* (Supplementary Fig. 2d)(28–30). These results indicate that macrophages in neuroblastomas have immunosuppressive features which may tune lymphoid responses.

**Figure 2.**
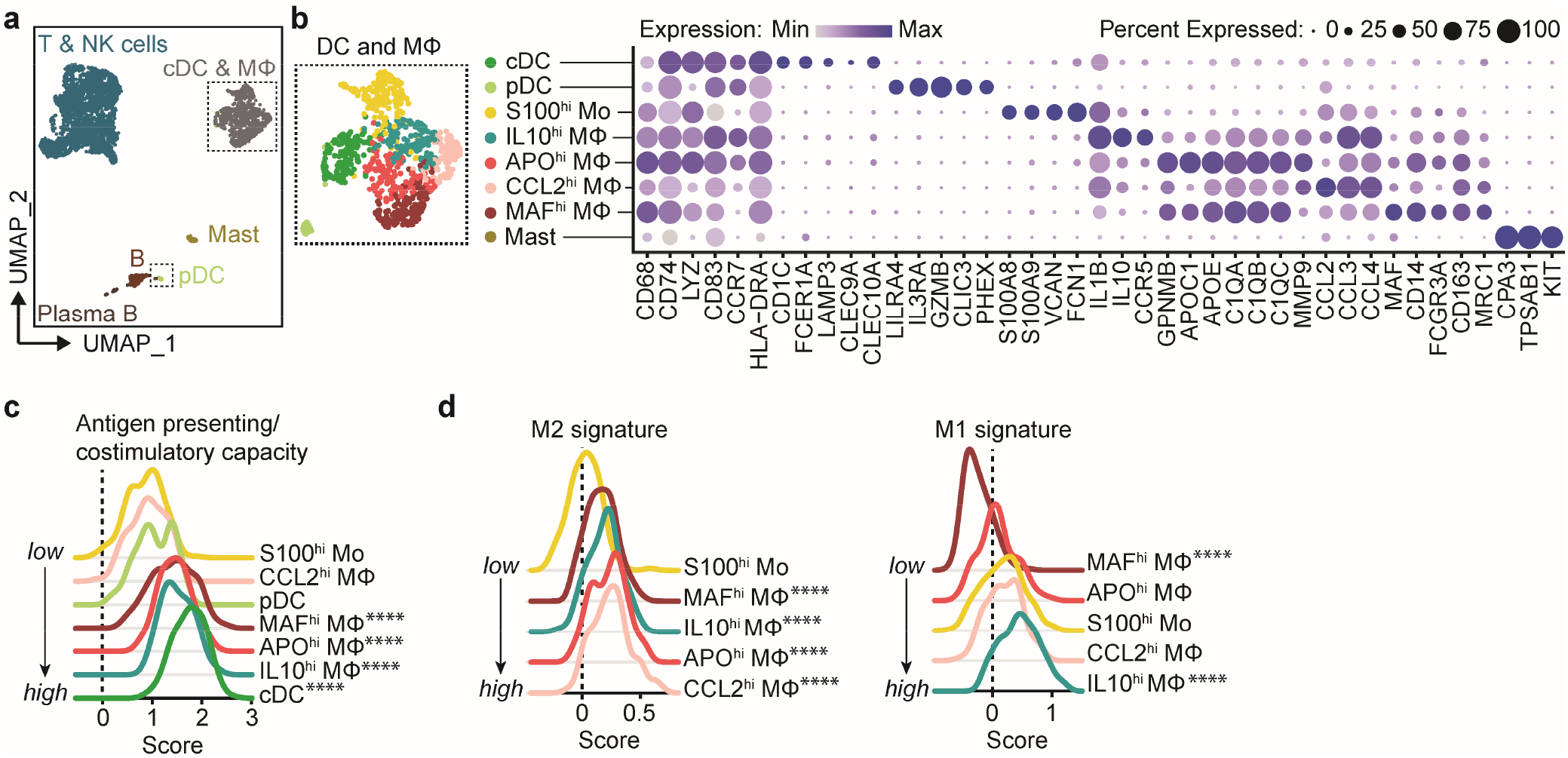
Neuroblastoma is populated by immunosuppressive macrophages. **(a)** UMAP of the immune compartment. Cells are colored by main immune cell types. **(b)** UMAP and dotplot of the myeloid compartment, showing conventional dendritic cell (cDC), plasmacytoid dendritic cell (pDC), undifferentiated monocyte (Mo) and differentiated macrophage (Mϕ) populations showing a selection of their marker genes. **(c)** Antigen presenting/co-stimulatory capacity score as constructed with genes in Supplementary Fig. 2c. **(d)** M1-like and M2-like macrophage signature expression score in monocytes and macrophages(26). *\*\*\*\*P<0*.*0001 versus S100*^*hi*^ *Mo, Kruskall Wallis with Dunn’s*.

### Lymphocyte subsets in neuroblastoma display degrees of dysfunctionality

The lymphoid compartment consisted of NK cells, T cells and a small fraction of (plasma) B cells (Fig. 2a and 3a). NK cells expressed genes encoding cytotoxic molecules (*GNLY, GZMA, GZMB, PRF1*), and γδ T cells expressed their specific T cell receptor (TCR) genes (*TRDC, TRGC1/2*). Among αβ T cells, we identified CD8^+^, CD4^+^, CD4^+^FOXP3^+^ regulatory T cells (Tregs), naive(-like) T cells expressing typical markers (*TCF7, SELL, LEF1, CCR7*), and a small heat-shock protein-positive cluster (HSP^+^ T; Fig. 3a and Supplementary Fig. 3a). CD8^+^ T cells expressed *CD8A* and cytotoxic effector molecules (*PRF1, GZMA, GZMB)*, and had significantly increased expression of *PDCD1* (encoding PD-1) and *LAG3* compared to other T/NK cell clusters (padj<0.0001; Fig. 3a,b). Since *PDCD1* and *LAG3* are typically regarded as markers of T cell dysfunction (also termed ‘exhaustion’), their expression indicated the presence of a dysfunctional cell fraction among CD8^+^ cells(31,32). Tregs expressed high levels of their signature genes *FOXP3* and *IL2RA*, in addition to transcription factors *IKZF2, BATF, PRDM1*, and *MAF* and checkpoint receptors *TNFRSF1B* (TNFR2), *TNFRSF4* (OX-40), and *TNFRSF18* (GITR) (Fig. 3c and Supplementary Fig 3b)(33–35). Expression of these transcription factors and checkpoint receptors is suggestive of an effector Treg profile, which is typically identified in tumor-infiltrating Tregs and associated with enhanced inhibitory capacity(36,37).

Among non-Treg CD4^+^ T cells we identified two distinct subsets with functional differences. One subset highly expressed *DUSP4*, which has been show to function as a signal repressor to prevent T cells from over-activation, and has been related to premature T cell aging causing senescence/exhaustion (Fig. 3d and Supplementary Fig. 3c)(38). This subset indeed showed a combined profile of activation and regulation, with TCR signaling and expression of *MKI67*, encoding proliferation marker Ki-67, on the one hand and high PD-1 signaling and regulatory genes (immune checkpoint receptors *TIGIT* and *CTLA4*, and transcription factors *TOX* and *TOX2)* on the other hand (Fig. 3a,d Supplementary Fig. 3c-f). Expression of these checkpoint receptors and transcription factors in non-Treg T cells has been previously associated with T cell dysfunction/exhaustion, thus suggesting the presence of dysfunctional cells among CD4^+^ T cells(31,39). Indeed, the *DUSP4*^hi^ CD4^+^ cluster showed significantly increased expression of a previously published signature for dysfunctional tumor-infiltrating CD4^+^ T cells (p<0.0001) (Fig. 3e)(32). The second cluster, containing *IL7R*^hi^ CD4^+^ T cells, had higher translational activity and expressed *CCR6* and *RORA*, both associated with Th17 polarization, and cytotoxic markers (*KLRB1, GNLY, PRF1*) (Supplementary Fig. 3c,d). These results suggest that the *DUSP4*^hi^ CD4 cells contained a higher proportion of tumor-reactive cells, whereas the *IL7R*^hi^ cells likely consisted of bystander cells. This premise was further supported by gene set enrichment analysis (GSEA) utilizing published gene signatures specific for either tumor-infiltrating T cells or blood/normal tissue-associated T cells(40), which showed that *DUSP4*^hi^ CD4 corresponded largely to previously identified tumor-infiltrating T cell clusters – including those with exhausted profiles – whereas *IL7R*^hi^ CD4 corresponded to blood/normal tissue-associated T cell clusters (Supplementary Fig. 3g). Taken together, the neuroblastoma immune environment is characterized by lymphocyte subsets with differing degrees of dysfunctionality, and highly immunosuppressive effector Tregs.

**Figure 3.**
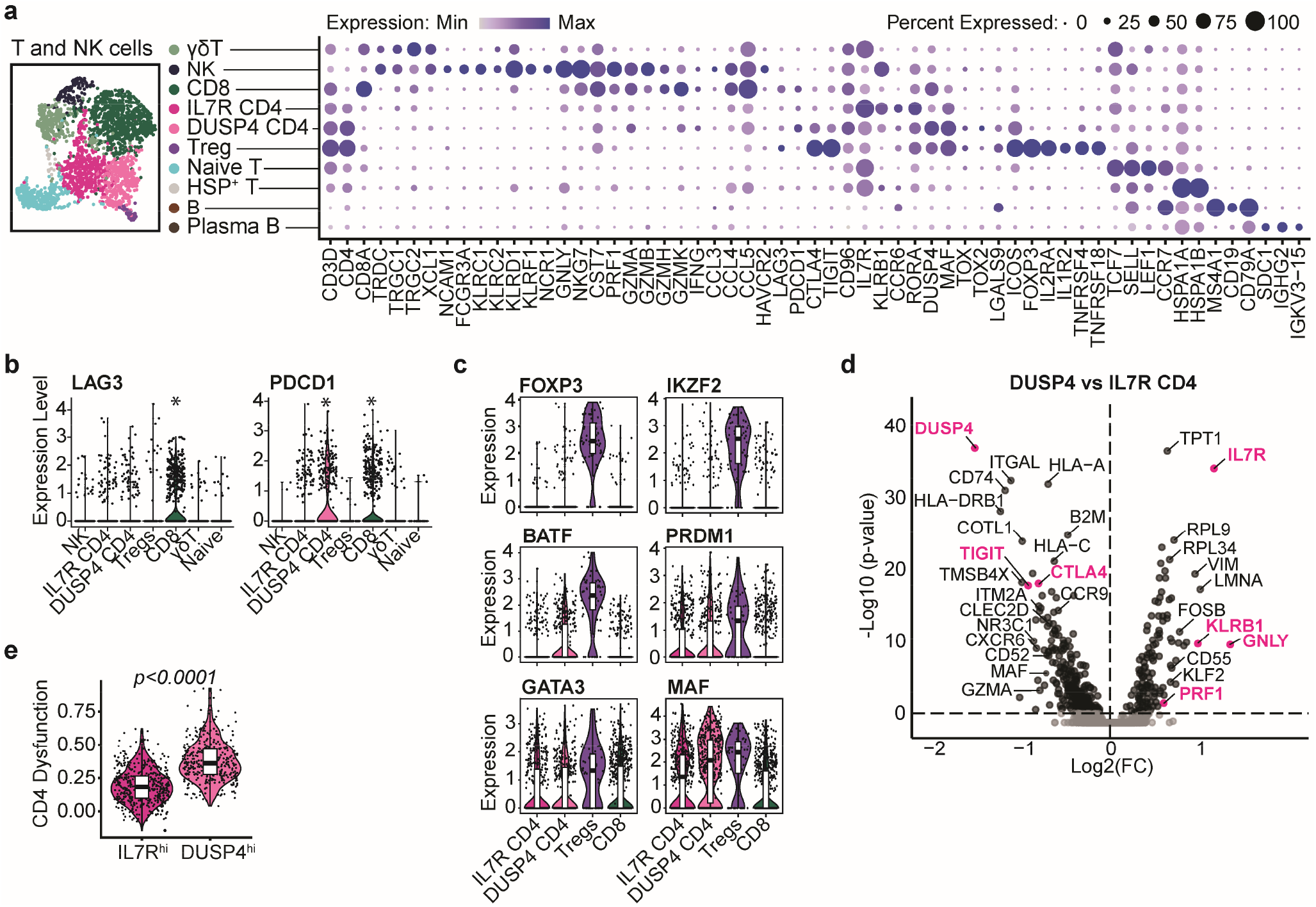
Neuroblastomas are characterized by lymphoid populations with differing degrees of dysfunctionality. **(a)** UMAP and dotplot of T and natural killer (NK) cell subclusters and a selection of their marker genes. **(b)** Expression of *LAG3* and *PDCD1* in T/NK cell clusters. \**Significantly upregulated with Padj<0*.*0001 in FindAllMarkers analysis*. **(c)** Transcription factors associated with effector Treg profile. **(d)** Volcano plot of differentially expressed genes between the two CD4 effector T cell populations. **(e)** Expression of a previously published signature for CD4 T cell dysfunction in melanoma(32).

### The immune cell composition and functional profile before and after induction chemotherapy

To assess the effect of induction chemotherapy on tumor immunity we compared the immune cell composition before (at diagnosis) and after induction chemotherapy (at surgical resection) (Fig. 4a). While the total number of immune cells was constant (Fig. 1f), the ratio of lymphoid to myeloid cells decreased upon treatment, suggesting either a reduction in lymphoid cells or an influx or expansion of myeloid cells (Fig. 4b and Supplementary Fig. 4a-c). Especially cDC and the strongly M2-differentiated *CCL2*^hi^ macrophages showed an increase (Fig. 4c and Supplementary Fig. 4d,e). In the lymphoid compartment, induction chemotherapy resulted in an effector-enriched milieu with decreased numbers of Tregs and naive cells, suggesting a shift towards a more immunogenic environment (Fig. 4d,e and Supplementary Fig. 4d,f). However, at the same time, NK cell numbers slightly decreased (Fig 4d and Supplementary Fig. 4d,f).

**Figure 4.**
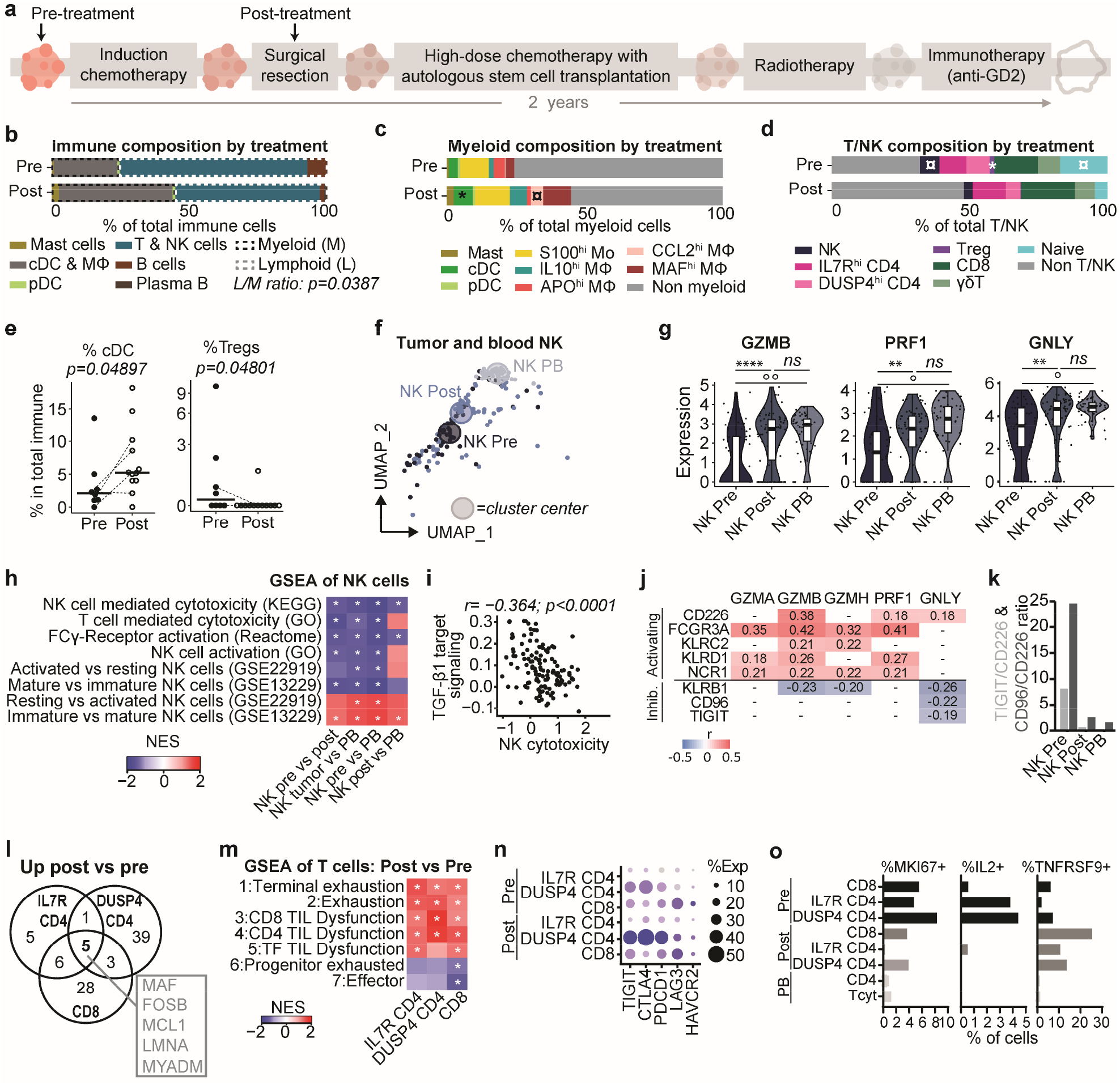
The immune cell composition and functional profile before and after treatment. **(a)** Schematic illustration of the high-risk neuroblastoma treatment plan. Arrows indicate sampling timepoints for single-cell RNA sequencing. **(b)** Average immune cell composition before and after induction chemotherapy. *Mann-Whitney U test*. **(c)** Average myeloid cell composition before and after induction chemotherapy. *\*significant difference (p<0*.*0*.*5) and ¤ trend (0*.*05<p<0*.*1) between pre and post*; *Mann-Whitney U test*. **(d)** Average lymphoid cell composition before and after treatment. *\*significant difference (p<0*.*0*.*5) and ¤ trend (0*.*05<p<0*.*1) between pre and post; Mann-Whitney U test*. **(e)** Percentage of cDC and Tregs of total immune cells before and after treatment. *Line-connected dots represent paired samples. Mann-Whitney U test*. **(f)** UMAP of NK cells from pre-treatment tumors, post-treatment tumors, and from healthy adult donor blood. Large circles indicate cluster centers. **(g)** Expression of cytotoxic genes by NK cells in tumors and peripheral blood (PB). *\*\*\*\*Nominal p<0*.*0001, ** Nominal p<0*.*01, °°padj<0*.*01, °padj<0*.*05* **(h)** GSEA of NK cells in pre-treatment tumors compared to NK cells from post-treatment tumors and blood. NK tumor = pre- and post-treatment NK cells combined. *\*FDR<0*.*25; NES=normalized enrichment score*. **(i)** Pearson correlation of a published TGF-β1 downstream signaling gene signature and NK cell cytotoxicity (modulescore of *GZMA, GZMB, PRF1, GNLY, NKG7, CST7, CCL5* and *IFNG*) in tumor NK cells (pre- and post-treatment combined)(43). **(j)** Pearson correlation coefficients of cytotoxic gene expression with expression of activating and inhibitory receptors in tumor-infiltrating NK cells. Significant correlations (p<0.05) are shown. ‘-’=not significant. **(k)** *TIGIT/CD226* and *CD96/CD226* gene expression ratios in NK cells from pre-/post-treatment tumors and from reference blood (PB). **(l)** Venn diagram of shared upregulated genes in CD4 and CD8 T cell subsets post-treatment compared to pre-treatment (padj<0.01). **(m)** GSEA of exhaustion and effector signatures (1+6: GSE84105, 2+7: Man et al.(49), 3+4+5: Li et al.(32)) in CD4 and CD8 T cells (\**FDR<0*.*25; NES = normalized enrichment score; TF=transcription factors*). **(n)** Dotplot of immune checkpoint receptor genes comparing expression in CD4 and CD8 T cells pre- and post-treatment. **(o)** Fraction of cells expressing proliferation marker *MKI67* (Ki-67), cytokine *IL2* (IL-2) and antigen-stimulated T cell marker *TNFRSF9* (4-1BB).

Since particularly NK cells are considered essential cytotoxic effectors in neuroblastoma as they are implicated in the efficacy of anti-GD2 therapy(41), we assessed whether their function changed upon treatment, and compared them to reference NK cells from healthy donor blood (Supplementary Fig. 5a,b). NK cells in pre-treatment tumors appeared to be more distinct from reference NK cells than NK cells in post-treatment tumors (Fig. 4f). As this could be due to micro-environmental effects induced by tumor cells, which were more abundant pre-treatment (Fig. 1f), we primarily focused on NK cell function pre-treatment. Compared to either NK cells post-treatment or reference NK cells (Supplementary Fig. 5c), NK cells in pre-treatment tumors had significantly reduced expression of essential cytotoxic effector genes (*GNLY, GZMB, PRF1*) (Fig. 4g), suggesting an impaired cytolytic function. GSEA confirmed their reduced cytotoxicity and indicated a more immature, resting state pre-treatment (Fig. 4h and Supplementary Fig. 5d), which has also been observed in other tumor types(42). Signaling by TGF-β1, a known immunosuppressive factor, was upregulated pre-treatment (Supplementary Fig. 5b) and correlated negatively with NK cell cytotoxicity (r=-0.364, p<0.0001; Fig. 4i)(43), indicating a possible role for TGF-β1 signaling in NK cell dysfunction. Since NK cell activity is additionally regulated by a delicate balance of activating and inhibitory receptors, we assessed expression of these receptors and evaluated their correlation with NK cell cytotoxicity (Supplementary Fig. 5e). Overall, activating receptor expression was lower in tumor NK than reference blood NK cells (Supplementary Fig. 5f). The activating receptors *CD226* (DNAM-1), *FCGR3A* (CD16), *KLRC2* (NKG2C), *KLRD1* (CD94) and *NCR1* (CD335) positively correlated with expression of cytotoxic genes, while the inhibitory *KLRB1* (CD161), *CD96* and *TIGIT* negatively correlated with cytotoxic gene expression (Fig. 4j). Of these, *KLRB1* correlated with 3/5 cytotoxic genes, implying broad effects on cytotoxicity. The activating *CD226* and inhibitory *CD96* and *TIGIT* belong to the same checkpoint receptor family competing for ligands. *TIGIT*/*CD226* and *CD96*/*CD226* gene expression ratios were substantially increased in tumor-infiltrating NK cells pre-treatment (Fig. 4k), indicating a disturbed balance shifted towards NK cell inhibition. These results implicate TGF-β1 signaling and the inhibitory checkpoint receptors KLRB1, TIGIT, and CD96 in the disturbance of NK cell cytotoxicity in neuroblastoma.

**Figure 5.**
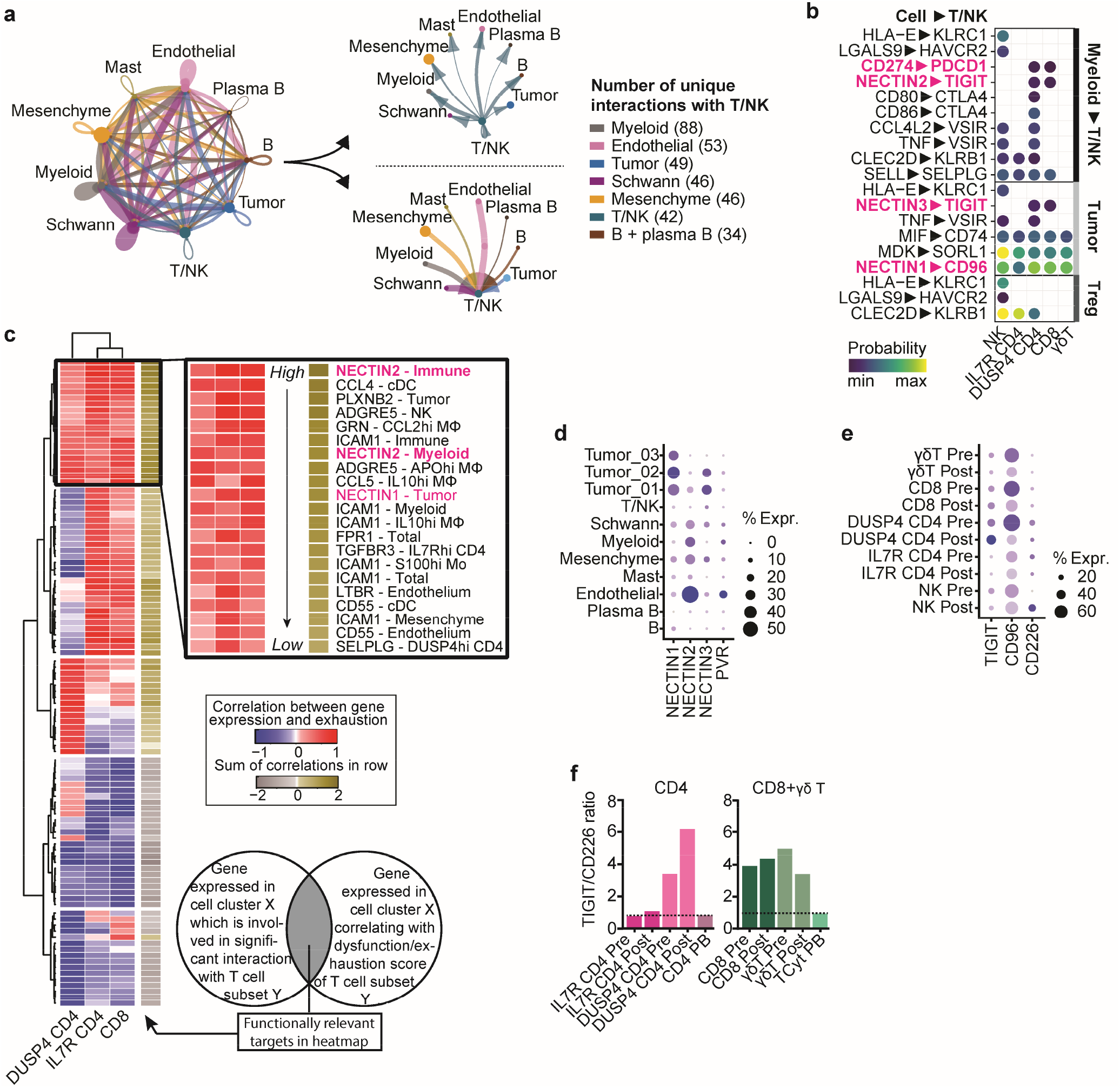
Immunoregulatory interactions are abundant in neuroblastoma. **(a)** Interaction network of main cell types in neuroblastoma constructed with CellChat(50). **(b)** Bubbleplot of selected, predicted immunosuppressive interactions between indicated T/NK subsets and myeloid cells, tumor cells and Tregs. Interactions with each specific myeloid subset were evaluated and subsequently merged, with the highest probability of each interaction pair depicted in the plot. For full overview see Supplementary Figure 7. **(c)** Heatmap showing genes which are involved in a significant ligand-receptor interaction between cell cluster X and an indicated T cell subset Y of which the expression by cell cluster X also significantly correlates with the exhaustion/dysfunction score of the indicated T cell subset Y. Genes with at least one significant correlation, with either *IL7R*^hi^ CD4, *DUSP4*^hi^ CD4 or CD8 T, were included. Black box indicates targets with the highest overall (sum) correlation with the dysfunction/exhaustion score. **(d**) Dotplot of *NECTIN1/2/3* and *PVR* expression in the neuroblastoma. **(e)** Dotplot of *TIGIT, CD96* and *CD226* expression in T/NK cells pre- and post-treatment. **(f)** *TIGIT/CD226* gene expression ratio in T cell subsets pre- and post-treatment and in reference blood.

### T cells in neuroblastoma show increased dysfunction post-treatment compared to pre-treatment

In contrast to NK cells, T cells did not have reduced cytotoxicity pre-treatment (Supplementary Fig. 6a). γδ T cells showed higher TCR activity and IL-12 and interferon signaling pre-treatment than post-treatment, suggesting that at least a fraction of γδ T cells in pre-treatment tumors was tumor-reactive (Supplementary Fig. 6b). αβ T cells (CD8^+^ and CD4^+^) showed similar results, also implying tumor-reactivity pre-treatment (Supplementary Fig. 6c,d). After treatment however, these T cell subsets shared a significantly increased expression of transcription factors *MAF, FOSB, MYADM, LMNA* and *MCL1* (Fig. 4l and Supplementary Fig. 6e). Of these, *MAF* and *MYADM* have been previously associated with exhaustion/dysfunction of T cells in tumor-microenvironments(32,44,45). GSEA confirmed that gene expression profiles previously linked to exhaustion and dysfunction were significantly enriched in post-treatment αβ T cells (FDR<0.25), whereas αβ T cells in pre-treatment samples had a higher effector signature (Fig. 4m). CD4^+^ and CD8^+^ T cells post-treatment were particularly enriched for signatures associated with ‘terminal exhaustion’ as opposed to ‘progenitor exhaustion’, the latter being associated with some level of retained functionality and tumor control(46). However, compared to healthy donor blood-derived T cells, both pre- and post-treatment tumor-infiltrating T cells showed a significant enrichment of progenitor and terminal exhaustion signatures (Supplementary Fig. 6f). The checkpoint receptors *LAG3* (LAG-3), *CTLA4* (CTLA-4), *PDCD1* (PD-1), *HAVCR2* (TIM-3), and *TIGIT*, which have been extensively linked to T cell exhaustion/dysfunction(31,32), were among the core enriched upregulated genes post-treatment (Fig. 4n and Supplementary Fig. 6g). Lower fractions of *MKI67* (Ki-67) and *IL2* (IL-2) expressing cells and increased fractions of *TNFRSF9* (4-1BB) expressing cells further confirmed reduced (proliferative) activity and suggested prolonged antigen stimulation (i.e. target recognition) post-treatment (Fig. 4o)(47,48). Taken together, these results imply that a fraction of T cells appears to be tumor-reactive and become more exhausted/dysfunctional after treatment, likely due to prolonged antigen stimulation. The high expression of immune checkpoints in combination with overall low TOX/TOX2 expression (Fig. 3a) however suggest potentially retained responsiveness to immune checkpoint blockade (ICB).

**Figure 6.**
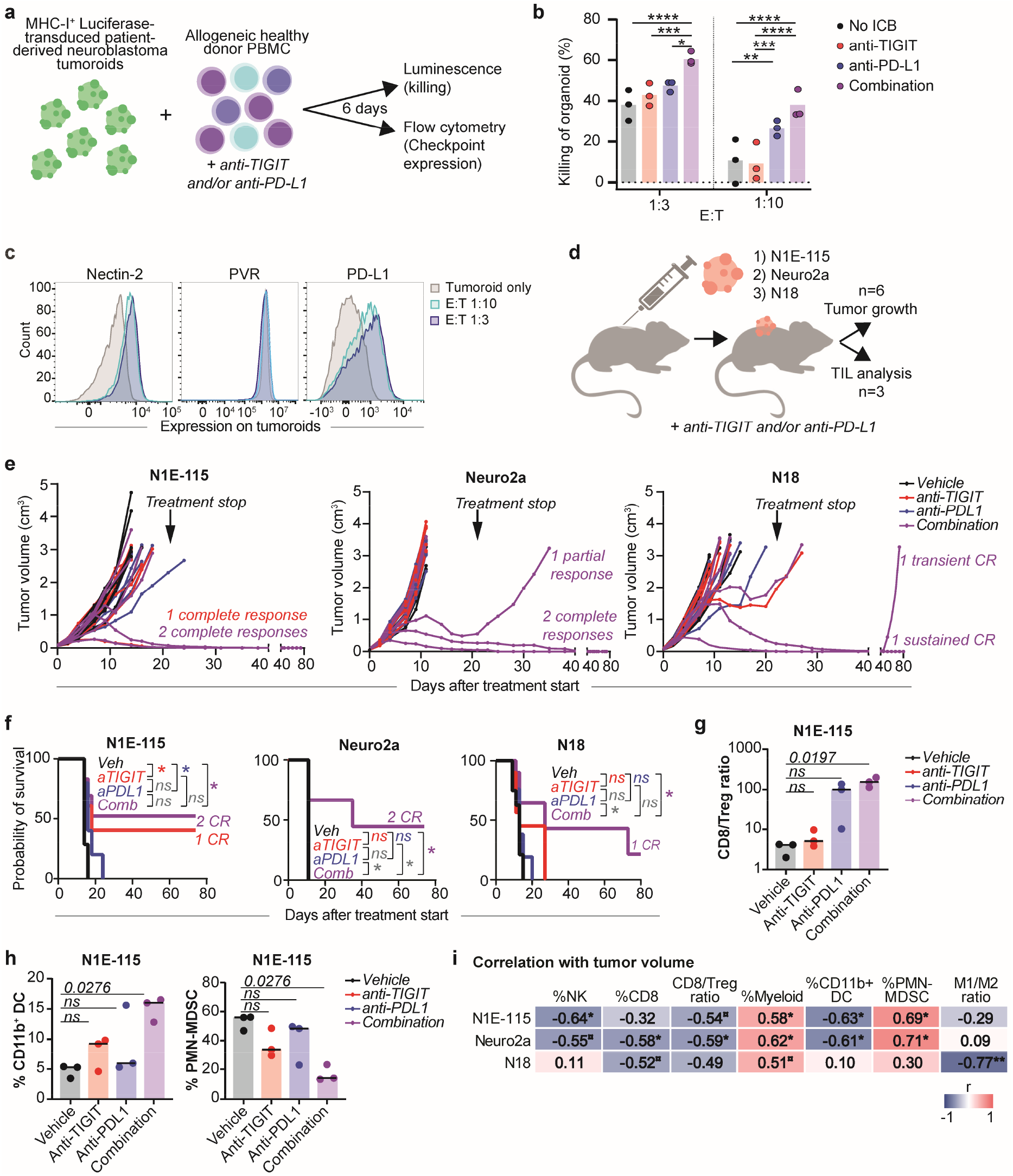
Combined TIGIT/PD-L1 blockade enhances immune responses against neuroblastoma. (**a**) Setup of in vitro killing assay with luciferase-transduced neuroblastoma tumoroids and healthy donor PBMC. Tumor cells and PBMC were cocultured for 6 days with or without anti-TIGIT and/or anti-PD-L1 at effector:target ratios 1:3 and 1:10 **(b)** Percentage of tumor cell killing (=100-normalized luminescence) of *in vitro* killing assay shown in Supplementary Fig.9b. Luminescence values were normalized against condition with untreated tumoroids only. *n=3. Two-way ANOVA with Tukey*.*\*p<0*.*05; **p<0*.*01, ***p<0*.*001, ****p<0*.*0001*. **(c)** Flow cytometric analysis of Nectin-2, PVR and PD-L1 expression on tumoroids cultured with or without PBMC at different E:T ratios. Gating strategy in Supplementary Fig. 9c. **(d)** Graphic representation of in vivo study. *TIL=tumor-infiltrating leukocytes*. **(e)** Tumor volume in N1E-115, Neuro2a and N18 mouse models (n=6 per group) treated with anti-TIGIT and/or anti-PD-L1 up to day 80 of follow-up. Treatment was discontinued after 3 weeks. **(f)** Survival analysis of N1E-115, Neuro2a and N18 models (n=6 per group) treated with anti-TIGIT and/or anti-PD-L1. Matched log-rank (Mantal-Cox) test indicated p values versus vehicle control. **(g+h)** Flow cytometric analysis of tumor-infiltrating leukocytes in N1E-115 model (n=3 per group) showing the CD8/Treg ratio **(g)** fraction of CD11b+ dendritic cells and polymorphonuclear myeloid-derived suppressor cells (PMN-MDSC) **(h)** in mice treated with anti-TIGIT and/or anti-PD-L1 for 7 days. *Kruskall-Wallis with Dunn’s*. **(i)** Correlation between tumor volume and tumor-infiltrating immune cell fractions in the three models (all four treatments combined). *Pearson correlation*.

### Immunoregulatory interactions in the neuroblastoma tumor-microenvironment

Lymphocyte dysfunction in tumors may result from prolonged antigen stimulation, but also from suppressive microenvironmental cues. To unravel these cues, we analyzed the interactions between T/NK cells and other cells in the tumor using the prediction tool CellChat(50). The combined population of T/NK cells (as defined in Fig. 2a) displayed a multitude of interactions with other cells, of which those with myeloid cells were most abundant (Fig. 5a; Supplementary Fig. 7a,b). To specifically identify therapeutically targetable immunomodulatory interactions, we focused on interactions between T/NK cells and three immunoregulatory interactions partners, i.e. myeloid cells, tumor cells and Tregs. Interactions with these three partners included, amongst others, interactions involved in cellular adhesion, chemoattraction and immune regulation (Supplementary Fig. 7a-e). Among these we identified the following cellular interactions with a potentially immunosuppressive role and potential for therapeutic intervention: *HLA-E—KLRC1* and *LGALS9—HAVCR2* for NK cells, *CD274—PDCD1, NECTIN2—TIGIT, CD80/CD86— CTLA4*, and *NECTIN3—TIGIT* for T cells, and *CCL4L2—VSIR, TNF—VSIR, CLEC2D—KLRB1, MIF—CD74, SELL—SELPLG, MDK—SORL1*, and *NECTIN1—CD96* for T and NK cells (Fig. 5b). These predicted cellular interactions clearly implicate immune checkpoint receptors CD161 (*KLRB1*), NKG2A (*KLRC1*), PD-1 (*PDCD1*), CTLA-4 (*CTLA4*), TIM-3 (*HAVCR2*), and the TIGIT/CD96 checkpoint receptor family in neuroblastoma. Moreover, the multitude of immunoregulatory interactions highlights a rationale for ICB combination therapy instead of ICB monotherapy.

**Figure 7.**
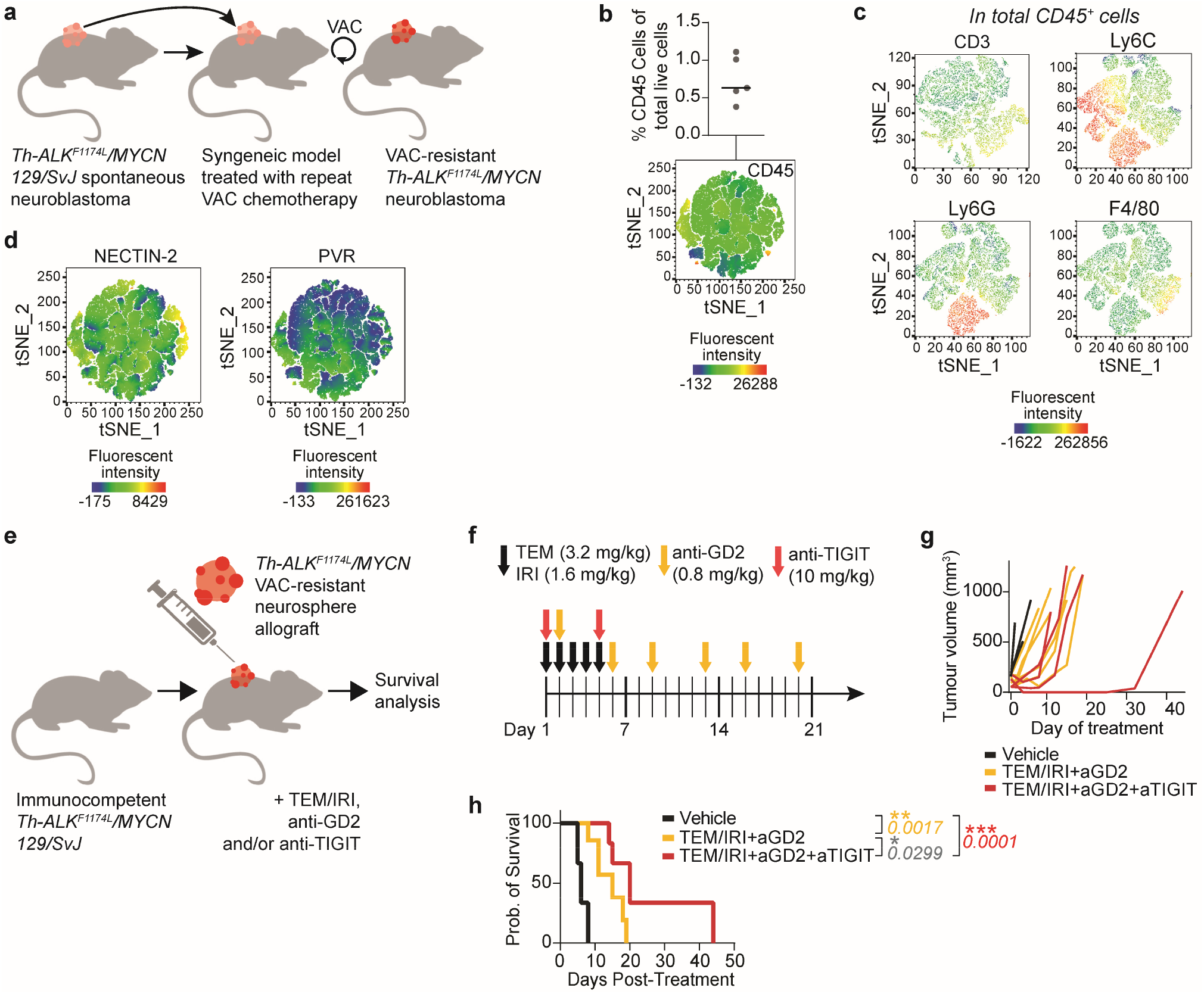
TIGIT blockade improves survival in a chemotherapy-resistant neuroblastoma model. **(a)** Generation of chemotherapy-resistant neuroblastoma model from *Th*-*ALK*^F1174L^/*MYCN* 129/SvJ models repeatedly treated with VAC (Vincristine, Adriamycin/Doxorubicin, Cyclophosphamide). **(b)** Percentage of CD45^+^ immune cells of total live cells in the tumor measured in tumor digest by flow cytometry. **(c)** Expression of CD3, Ly6c, Ly6G and F4/80 among CD45^+^ tumor-infiltrating immune cells measured in tumor digest by flow cytometry. (**d**) Expression of Nectin-2 and PVR among total live cells in tumor digest measured by flow cytometry. (**e**) Schematic representation of study setup adding anti-TIGIT to the standard relapse backbone treatment (Temozolomide/Irinotecan (TEM/IRI) + anti-GD2). **(f)** Treatment schedule for TEM, IRI, anti-GD2 and anti-TIGIT. **(g)** Tumor volume measured over time in mice with vehicle (n=3), TEM/IRI + anti-GD2 (n=6) or TEM/IRI + anti-GD2 + anti-TIGIT (n=4) treatment. **(h)** Survival analysis in mice with vehicle (n=3), TEM/IRI + anti-GD2 (n=6) or TEM/IRI + anti-GD2 + anti-TIGIT (n=4) treatment. *Matched log-rank (Mantel-Cox) test*.

### Immunosuppressive interactions with potential for therapeutic intervention

Immune checkpoint blockade has revolutionized cancer therapy and is thought to restore activity of exhausted/dysfunctional T cells(1). We therefore sought to identify more specifically which of the interactions between T cells and other cells were associated with disturbed T cell function, with the goal of therapeutically reinvigorating T cell function by targeting exactly those interactions. We assessed the correlation between αβ T cell dysfunction and predicted interactions with an unbiased approach, as outlined in Fig. 5c, using a dysfunction/exhaustion score with established markers (Supplementary Fig. 8a,b)(51). The subsequent interaction analysis revealed 13 unique target interactions which positively correlated with dysfunction/exhaustion of αβ T cells (Fig. 5c and Supplementary Fig. 8c). The top target, showing the highest (sum of) correlations with T cell dysfunctionality, was *NECTIN2* expressed by immune cells. Although *NECTIN2* was mostly expressed by endothelial cells (Fig. 5d), specifically *NECTIN2* expressed by cDC, *IL10*^hi^ Mϕ and *APO*^hi^ Mϕ was predicted to interact with *TIGIT* on CD8 and *DUSP4*^hi^ CD4 T cells (Supplementary Fig. 8c). We confirmed the correlation between *NECTIN2* expression and exhaustion scores in bulk-RNAseq data of 498 neuroblastomas (SEQC cohort GSE49710; Supplementary Fig 8e)(52). The other NECTIN-family members *NECTIN1* and *NECTIN3*, which compete for interactions with *NECTIN2*, were predominantly expressed by tumor cells (Fig. 5d) and predicted to interact with *CD96* on T and NK cells and *TIGIT* on T cells, respectively (Fig. 5b and Supplementary Fig. 8c). *PVR* was expressed by endothelial cells but did not have significantly predicted interactions (Fig. 5d). Like in neuroblastoma-infiltrating NK cells (Fig. 4k), the balance between NECTIN-binding inhibitory receptors *TIGIT* and *CD96* and activating receptor *CD226* proved disturbed in *DUSP4*^hi^ CD4, CD8 and γδ T cells when compared to reference T cells from blood (Fig. 5e,f and Supplementary Fig. 8f). Taken together, NECTIN family members may regulate T/NK cell function in neuroblastoma, and particularly the *NECTIN2—TIGIT* interaction may represent a promising target for therapeutic intervention.

### Combined TIGIT/PD-L1 blockade enhances immune responses against neuroblastoma

To investigate the therapeutic potential of TIGIT blockade in neuroblastoma, we explored its functional role in tumor killing. To consider the possible benefit of ICB combination therapy – based on the abundance of other immunoregulatory interactions (Fig. 5b) – we also investigated combined TIGIT/PD-L1 blockade. PD-L1 was highly expressed by myeloid cells (Supplementary Fig. 9a). TIGIT blockade *in vitro* resulted in significantly enhanced killing of neuroblastoma tumoroids by immune cells, when combined with PD-L1 blockade (Fig. 6a,b and Supplementary Fig. 9b-d). The rationale for this ICB combination was further substantiated by upregulation of TIGIT and PD-1 ligands Nectin-2 and PD-L1 by tumor cells during coculture (Fig. 6c). *In vivo*, combined TIGIT/PD-L1 blockade led to sustained complete remissions (CR) in 2 out of 6 animals in two murine models (N1E-115 and Neuro2a) in addition to a partial response in Neuro2a, and 1 transient CR and 1 sustained CR in a third model (N18) (Fig. 6d-f and Supplementary Fig. 10a). None of the treatments led to changes in body weight or other toxicities (Supplementary Fig. 10b). The combined treatment significantly modulated the immune environment into a more immune-activated state, with an increased CD8/Treg ratio, increased CD11b^+^ DC numbers and reduced immunosuppressive polymorphonuclear myeloid-derived suppressor cell (PMN-MDSC) numbers in the N1E-115 model (Fig. 6g,h and Supplementary Fig. 10c). In Neuro2a and N18 mice, similar trends were observed (Supplementary Fig. 10c,d). Percentages of NK, CD8, CD11b+ DC as well as CD8/Treg and M1/M2 ratios negatively correlated with tumor volume in the mouse models, suggesting a role in tumor eradication (Fig. 6i). In contrast, overall myeloid and PMN-MDSC percentages positively correlated with tumor volume, confirming their pro-tumorigenic potential. In conclusion, we identified TIGIT as a relevant immune checkpoint in neuroblastoma, with therapeutic potential in combination with PD-L1 blockade.

### TIGIT blockade improves survival in a chemotherapy-resistant neuroblastoma model

Since tumor relapses are the major cause of the low survival rates in patients with neuroblastoma, and novel treatments are especially urgent for this group of patients, we next investigated the added effect of TIGIT blockade in an immunologically cold, chemotherapy-resistant syngeneic model. The model was generated by allograft of neurospheres derived from *Th*-*ALK*^F1174L^/*MYCN* 129/SvJ transgenic spontaneous tumors subjected to repeat chemotherapy (Fig. 7a and supplementary figure 11a-c)(53). CD45^+^ immune cells only made up ∼1% of total live cells, but included CD4, CD8 and γδ T cells and Ly6G^+^ PMN-MDSC and Ly6C^+^ or F4/80^+^ monocytes/Mϕ (Fig. 7b,c and supplementary figure 11d). Since the TIGIT ligands NECTIN2 and PVR were also expressed in this model, we next investigated the potential of TIGIT blockade (Fig. 7d). TIGIT blockade was added to temozolomide and irinotecan (TEM/IRI) + anti-GD2 antibody treatment, which is favored as the standard backbone treatment for relapse/refractory neuroblastoma patients in Europe (Fig. 7e,f). Addition of TIGIT blockade to TEM/IRI+anti-GD2 significantly improved animal survival compared to TEM/IRI+anti-GD2 alone (p=0.0299) and compared to the vehicle control (p=0.0001; Fig. 7g,h). TIGIT blockade may thus add a significant survival benefit to the currently favored relapse treatment protocol, even in the context of chemotherapy resistance, and has high potential for translation into the clinical (relapse) setting.

## DISCUSSION

In this study we have generated a comprehensive single-cell atlas of the neuroblastoma immune environment, revealing the detailed composition and functional profile of immune cells in neuroblastoma. We exposed a vast immunoregulatory network affecting T and NK cell function and identified TGF-β1, CD161 (*KLRB1*), NKG2A (*KLRC1*), PD-1 (*PDCD1*), CTLA-4 (*CTLA4*), TIM-3 (*HAVCR2*), CD96 and, above all, TIGIT as promising targets for immunotherapy. *In vitro* and *in vivo* studies validated combined TIGIT/PD-L1 blockade as a novel, viable treatment option for patients with neuroblastoma. cells in pre-treatment tumors – suggests that immunotherapy may be able to reverse or at least ameliorate the arrested state of NK cells(54), as also previously observed in peripheral blood of patients.(55). Still, NK cell dysfunctionality in neuroblastoma may warrant further investigations into parent-derived NK cell infusions in combination with anti-GD2.

Our analyses revealed that effector lymphocytes in neuroblastoma are dysfunctional, as also observed in other tumors(8,9,40). NK cells in neuroblastoma had reduced cytotoxicity, particularly pre-chemotherapy. This resting, immature profile of NK cells recapitulates their previously described state in other tumors, designated as ‘arrested development’(42). NK cell dysfunction in neuroblastoma is not limited to the tumor microenvironment but has also been observed in peripheral blood of patients, throughout the treatment course(55). A recently published phase II trial adding anti-GD2 treatment, which is considered to rely at least partly on activity of NK cells(41), to conventional induction chemotherapy, observed improved early responses and event-free survival in comparison with historic reference cohorts(54). This added value of anti-GD2 during the induction phase – despite the dysfunctionality NK While T cells in neuroblastoma showed signs of tumor reactivity pre-treatment, they exhibited features of dysfunction post-treatment, characterized by high levels of immune checkpoint genes *LAG3, PDCD1, TIGIT, CTLA4* and *HAVCR2*(8,56). T cell dysfunction may result from chronic stimulation of T cells and/or immunomodulatory signals(1,57). Considering the high expression of antigen-experienced T cell marker *TNFRSF9* (4-1BB) post-treatment, prolonged antigen exposure likely contributed to the development of T cell dysfunction(47,48); months of induction chemotherapy may have increased tumor cell immunogenicity, eliciting chronic T cell activation with subsequent exhaustion(58). However, we cannot rule out that chemotherapy may have also directly contributed to induction of dysfunctional features in T cells post-treatment(59). Yet, the high expression of immune checkpoints in combination with overall low TOX/TOX2 expression suggests potential retained responsiveness to ICB.

We revealed an abundance of immunomodulatory interactions, involving immune checkpoints CD161, NKG2A, PD-1, CTLA-4, TIM-3, CD96 and TIGIT, between T/NK cells and three interactions partners tumor, effector-differentiated Tregs and suppressive macrophages. Effector-differentiation of Tregs, observed in various tumors, has been related to enhanced suppressive capacity(36,37). Neuroblastoma was dominated by M2-like-differentiated macrophages, associated with regulatory roles and tumorigenic potential(26,27). These have recently moved into the spotlight as targets for immunotherapy: combined anti-GD2/CD47 therapy to activate Mϕ resulted in highly promising anti-tumor activity, accompanied by recruitment of M1-like Mϕ and reduced M2-like Mϕ(60). Taken together, the abundance of T/NK cell interactions with highly immunosuppressive cells warrants exploration of immunotherapy combination strategies.

We identified the immunoregulatory *NECTIN2—TIGIT* axis as a target with high potential for therapeutic intervention. *NECTIN2—TIGIT* interaction has been reported in other solid tumors(61,62), underlining its universality in tumor microenvironments. Blockade of TIGIT has been shown to not only increase T cell, but also NK cell antitumor immunity(63,64). Moreover, combined TIGIT/PD-L1 blockade enhanced antitumor responses in other solid cancers *in vitro* and *in vivo* and the observed *in vivo* responses shared high similarity with ours, resulting in complete responses in a part of the animals(63,64). This heterogeneous response to ICB in syngeneic animal models has been previously related to Mϕ-driven ICB resistance(65), which suggests the potential of therapeutic interventions directed at Mϕ to increase response to ICB and highlights the importance of identifying biomarkers for treatment response. Since the patient numbers in our study were too small to perform patient subgroup analyses, this would be a valuable additional analysis in future studies Importantly, the first clinical trials combining TIGIT and PD-1/PD-L1 blockade have produced encouraging results in advanced solid tumors in adults(66–68).

As an ultimate effort of translation to an early phase clinical trial, by mimicking the immune environment of pre-treated patients with chemotherapy-resistant tumors, we demonstrated a survival benefit of adding TIGIT blockade to the currently favored relapse backbone treatment in a chemotherapy-resistant, immunologically cold, syngeneic mouse model with shared embryological origins to the human counterpart. These findings open the opportunity for clinical development of alternative checkpoint inhibitors such as anti-TIGIT in (pediatric) solid cancers. Moreover, the vast immunoregulatory network in neuroblastoma provides a rationale for immunotherapy combination strategies. The number and complexity of the immunosuppressive signaling axes offers a likely explanation for the lacking efficacy of immunotherapy with a single agent(18,19). To overcome immunosuppressive signaling and reinvigorate T/NK cell responses, a combination of immunotherapies targeting multiple immunosuppressive pathways at once, may be required. The already described anti-GD2/CD47 combination, and a case report of two refractory neuroblastoma patients treated with anti-GD2/PD-1 combination therapy illustrate the potential of such combinations(60,69). Importantly, future clinical studies investigating combination strategies will have to balance immunotherapy efficacy with the risk of immunotoxicity, and consider biomarkers to predict treatment response. In conclusion, we have constructed a comprehensive atlas of the neuroblastoma immune environment and identified functionally relevant targets, including TIGIT, for novel (combination) immunotherapies.

## METHODS

### Ethics statement

Studies underlying this paper have received appropriate approval by ethics review boards as per national legislation. The studies were conducted in patients in accordance with the Declaration of Helsinki. Age-appropriate written informed consent from patients and/or parents was obtained prior to inclusion of each patient in the study. Dutch tumor samples were obtained through an institutionally approved research study by the biobank committee of the Princess Máxima Center.

### Participants

From the Princess Máxima Center in Utrecht, The Netherlands, 20 patients with neuroblastoma were included between September 2017 and November 2020. Samples were obtained from patients during diagnostic biopsy pre-treatment (n=10) or during surgical resection after ca. 5 months of chemotherapeutic treatment (n=15). Demographic (e.g. sex and age), clinical (e.g. treatment) and histological information, including tumor staging by INRG, INRGSS and INSS was collected. The presence of MYCN amplification was determined by fluorescence in situ hybridization at the pathology department. 5 adult healthy volunteers donated blood for analysis of healthy immune cells.

### Isolation of cells from tumor biopsies

Tumor material was collected in the operating room by Truecut biopsy (samples at diagnosis) or surgical resection (debulking procedure after induction chemotherapy). The Dutch samples were gently dissociated into single-cells and sorted by flow cytometry for single-cell RNA sequencing using the CEL-Seq2 platform. In brief, preparation of the tumor pieces for single-cell RNA sequencing was started within 4 hours after the surgery. The material was minced into pieces <1mm^3^ and dissolved by enzymatic digestion with collagenase I, II and IV (2.5 mg/mL) at 37°C with agitation for max. 1 hour. The samples were filtered through a 70 μm cell strainer and washed in DMEM. Cells were further separated into a single cell suspension with the NeuroCult dissociation kit according to the manufacturer’s protocol (Stemcell™ Technologies, cat#05707), washed and stained with DAPI and DRAQ5. Next to this unbiased strategy, some samples were additionally stained with antibodies to enrich for T cell populations (NB124, NB125, NB138 with CD3-PE (Biolegend, cat#300308); NB124 with TCRγδ-BV421 (Biolegend, cat#331217) or with GD2-FITC (BD Biosciences, cat#563439) to enrich for tumor cells (000GGU, 000GXF, NB106, NB107, NB098, NB125, NB125, NB138, NB130, NB152). Single live cells were sorted into 384-well plates containing 10 μl of mineral oil, 50 nl of RT primers, deoxynucleotide triphosphates (dNTPs) and synthetic mRNA Spike-Ins on a FACSJazz, FACSAria II or Sony SH800S machine and subsequently spun down, snap-frozen on dry ice, and stored in -80°C to proceed with the total transcriptome amplification, library preparation and sequencing.

### Isolation of cells from donor peripheral blood

Peripheral blood mononuclear cells (PBMC) from 5 healthy young adult donors (mean age 28) were isolated by Ficoll-Paque (VWR) density centrifugation and stained with fixable viability dye efluor506 (eBioscience, cat#65-0866-14), CD3-PE (Biolegend, cat#300308) and CD56-APC (Biolegend, cat#318310) antibodies. Subsequently, live CD3^+^ T cells and CD56^+^ NK cells were FACS sorted into 384-well plates containing 10 μl of mineral oil, 50 nl of RT primers, deoxynucleotide triphosphates (dNTPs) and synthetic mRNA Spike-Ins on a Sony SH800S machine. The plates were spun down, snap-frozen on dry ice, and stored in -80°C to proceed with the total transcriptome amplification, library preparation and sequencing.

### CEL-Seq2 library preparation, sequencing, and mapping

All samples were processed for total transcriptome amplification, library preparation and sequencing into Illumina sequencing libraries as previously described(70). Paired-end 2×75 bp sequencing read length was used to sequence the prepared libraries using the Illumina NextSeq sequencer. Sharq preprocessing and QC pipeline were applied to process the single-cell RNA-seq data as described(71). Read mapping was done using STAR version 2.6.1 (RRID:SCR_004463) on the hg38 Patch 10 human genome. Function featureCounts (RRID:SCR_012919) of the subread package (version 1.5.2) was used to assign reads based on GENCODE version 26 (RRID:SCR_014966).

### CEL-Seq2 quality control and downstream analysis of single-cell RNA-seq data

Failed reactions were identified by low levels of ERCC external RNA controls and excluded(71). Furthermore, a liveness threshold was calculated for each plate based on the wells with no cell added, in order to distinguish live cells from dead and/or apoptotic cells(71). This threshold was set to a minimum of 500 transcripts. Next, the percentage of transcripts mapping to the mitochondrial genome was calculated and cells with more mitochondrial-encoded transcripts over nuclear ones were removed. Mitochondrial and ERCC transcripts were removed from the dataset, as well as cells with <1000 nuclear-encoded transcripts or <500 genes. In addition, cells with >150.000 nuclear-encoded transcripts were removed. Genes with low expression, that is either having less than 5 cells expressing the gene or less than two cells with less than two transcripts, were removed. A total of 105 cells, forming a distinct cluster of erythroid lineages, was identified based on high levels of hemoglobin complex genes, and was removed. To improve cross-sample comparisons, ambient mRNA contamination in individual cells was estimated and removed using DecontX(72). DecontX was run for all samples (batches) individually. Removal of cells with less than 1000 nuclear-encoded transcripts was repeated on the decontaminated counts matrix outputted by DecontX.

All subsequent analyses were performed using R (version 4.0.2) and the package Seurat (version 3.2.2, RRID:SCR_007322) with default parameters unless stated otherwise. The *SCTransform*() function was used to normalize and scale the data, and to identify variable genes. To avoid clustering of cells based on specific cell processes, genes associated with sex (*XIST, TSIX*, and Y chromosome-specific genes), cell cycle phase, dissociation stress (heat shock proteins; GO:0006986), and activity (ribosomal protein genes; GO:0022626), were removed as described before(73).

Principal component analyses were performed using the filtered lists of variable genes. To study the main cell types the first 35 principal components (PCs) were used to calculate dimensionality reduction using UMAP, and a resolution of 0.5 was used to define clusters using the Louvain method. For immune cell-focused analysis immune cell clusters were subset based on PTPRC gene expression. Clustering was performed using the same principal components and a resolution of 0.3 was used to define clusters of the main immune cell types. For in-depth analysis of the T and NK cells the respective clusters were subsetted, UMAP was rerun using 40 PCs and a resolution of 0.7 was used to define subclusters. For in-depth analysis of the myeloid compartment, UMAP was rerun using 35 PCs and a resolution of 0.8 was used to define subclusters.

### Cluster annotation

Cluster annotation was performed using the R package SingleR (version 1.2.4)(74), using the HumanPrimaryCellAtlas reference dataset to annotate main cell types, and additionally using the *NovershternHematopoieticData*(75) and *MonacoImmuneData*(76) reference datasets to annotate the immune cell (sub)clusters. Cell annotations were further refined by consulting cluster-specific (up-regulated) differentially expressed marker genes using Seurat’s FindAllMarkers function. The outputted genes were compared to known cell-type specific marker genes from previous studies(32,77–80) Malignant and non-malignant cells were distinguished according to three criteria: (1) their inferred CNV profiles (*see below*)(81); (2) under-expression or absence of different non-malignant cell type marker genes; and (3) high expression of published neuroblastoma-associated genes (Fig. 1d and Supplementary Fig. 1b). For tumor cell identification by copy number variation inference the R package inferCNV was run (with default settings except cutoff=0.1 and denoise=T) using the immune cell clusters as a reference.

### Differential gene expression analysis

Cluster-specific genes were identified using the *Findallmarkers* function in Seurat (RRID:SCR_007322) and genes with a padj<0.05 were considered differentially expressed. Differentially expressed genes between two groups were identified using the *Findmarkers* function and genes with a padj<0.1 or p<0.05 were considered differentially expressed, as indicated in the figure legends. Volcanoplots were created with the R package EnhancedVolcano.

### Cellular composition analysis

For analyses determining the composition of cell types of the individual tumor samples, we only included samples which were sorted in an unbiased manner (DAPI & DRAQ5 for total cells, overall immune cells, and myeloid cells; DAPI & DRAQ5 or anti-CD3 for T cell subsets). For immune cell composition analyses, samples with <10 immune cells were excluded. Barplots were generated using ggplot2 (version 3.3.2, RRID:SCR_014601) and visualize the percentage of cells per neuroblastoma sample. Differences in composition between treatment-naive and treatment-exposed tumor samples were statistically tested using the Wilcoxon rank-sum test.

### Pathway and gene set enrichment analysis

Pathway enrichment analysis was conducted using publicly available online Reactome portal (https://reactome.org), and pathways with >10 identified genes and Bonferroni-corrected *P*<0.05 were considered statistically significant. For GSEA, differential expression analysis between two groups was performed with *Findmarkers* using adjusted parameters (see code). Genes were pre-ranked by their Fold Change and GSEA was performed using Broad Institute software, by 1000 random permutations of the phenotypic subgroups to establish a null distribution of enrichment score, against which a normalized enrichment scores and multiple testing FDR-corrected *q* values were calculated(82). Gene sets with an FDR<0.25 were considered significantly enriched, as recommend by Broad Institute. Gene sets were either obtained from provided data in publications or by analyzing raw data using GEO2R (NCBI tool)(83). An overview of used signatures is provided in Supplementary Table 2.

### Interaction analysis

The CellChat algorithm was applied to perform an unbiased ligand-receptor interaction analysis, using the curated ligand-receptor database of CellPhoneDB (RRID:SCR_017054)http://www.cellphonedb.org/(50). To identify functionally relevant interactions, we overlaid 1) genes expressed by cell subset X which were predicted to be membrane-expressed or secreted in the Human Protein Atlas database(84) and had a significant correlation (p<0.05) with the cytotoxicity score or exhaustion score of T/NK cell subset Y and 2) were predicted to have a significant interaction (p<0.05) between cell subset X and T/NK cell subset Y.

### In vitro tumor killing assay with immune checkpoint inhibitors

Patient-derived tumoroids(AMC691T: GD2^-^ MHC-1^+^) were transduced with GFP and an endogenous luciferase construct as described before(85). A single-cell solution was prepared using enzymatic (Accutase, Sigma) and mechanical dissociation. 5000 cells per well were seeded in 100 μl tumoroid medium in a white flat-bottom TC-treated 96-well plate (Corning) and rested for 3 days to reform spheres at 37°C and 5% CO2. Effector cells (isolated allogeneic PBMCs from healthy donors) were added to the tumoroids in a 1:3 and 1:10 effector:target (E:T) ratio in RPMI 1640 (Gibco) supplemented with 10% FBS, 100 U/mL Penicillin/Streptomycin and 2mM% L-Glutamine, with or without anti-TIGIT (clone 10A7) and/or anti-PD-L1 (clone 6E11; 10 ng/mL). Both antibodies were kindly provided by Roche. After 6 days, D-luciferin (150 μg/mL; Perkin Elmer) was added to the culture and incubated for 5 minutes at 37°C. The produced luminescence signal was detected with the FLUOstar Omega microplate reader. This signal was used as a readout measure for viable cells. A tumoroid only control was used to normalize and calculate the killing capacity.

### In vivo experiments with immune checkpoint inhibitors

In vivo experiments were carried out in Neuro-2a, N1E-115 and N18 (Hamprecht) syngeneic tumor models. Approximately eight-week-old female A/Jax-000646 mice were ordered from The Jackson Laboratory. Neuroblastoma cells were routinely subcultured twice weekly. The cells growing in an exponential growth were subcutaneously inoculated at 1×10^6^ cells/ml in 0.1 ml PBS for tumor development. On day 0, when average tumor volume was ∼100 mm3, body weight and tumor volume were measured, and mice were randomized into study groups with nine animals per arm. Mice were dosed IP with anti-mu-TIGIT (10 mg/kg, first dosing was done i.v.) and anti-PDL1 clone 6E11 (10mg/kg), supplied by Roche. Both molecules were administrated 3 times per week for 3 weeks. For TIL analysis tumors were harvested, weighed, and processed to obtain single-cell suspensions; tumor tissues were dissociated using MACS Dissociator (Miltenyi-130-093-235). All samples were analyzed by flow cytometry (BD FACS LSR Fortessa) and >10,000 CD45+ cells were recorded for further analysis.

### Immune checkpoint inhibition in a chemotherapy-resistant in vivo model

The chemotherapy-resistant model was generated by allograft of neurospheres derived from *Th*-*ALK*^F1174L^/*MYCN* 129/SvJ transgenic spontaneous tumors into syngeneic animals, which were then subjected to repeat and escalating chemotherapy (VAC – Vincristine, Adriamycin/Doxorubicin, Cyclophosphamide). These chemotherapy-resistant tumors were subsequently allografted into untreated syngeneic animals which underwent treatment with TEM/IRI (Temozolomide: 2.3 mg/kg IP; Irinotecan: 1.6 mg/kg IP, both days 1-5) + anti-GD2 (14G2a, 0.8 mg/kg IP, day 1 and 5) with or without anti-TIGIT (10mg/kg IP, twice weekly every week until termination). Animals with tumors >200mm^3^ before start of treatment were excluded from the analysis.

### General statistics

Apart from the differential gene expression analysis described above, comparisons between two groups (e.g. for signatures) were made by Mann-Whitney U test. Comparisons between 3 or more groups were analyzed by Kruskall-Wallis with Dunn’s post hoc test for multiple comparisons. For heatmap analysis, normalized gene expression was extracted from DotPlot analysis in Seurat, and hierarchical clustering was performed with Ward’s method and Euclidian distance. For correlation analyses, Pearson correlation coefficients were computed. FACS data were analyzed using FlowJo V10.8.1 software (LLC, RRID:SCR_008520). Statistical analysis of FACS data was performed with Graphpad Prism (GraphPad Software, RRID:SCR_002798). For survival analysis, matched analysis using the log-rank (Mantel-Cox) test was performed.

### Data access statement

The code and single-cell RNA sequencing data will be made available upon reasonable request with the authors.

## Acknowledgements

We would like to acknowledge the head of our FACS facility Tomasz Poplonski for his help and support. This work has received funding from the European Union’s Horizon 2020 research and innovation program under the Marie Skłodowska-Curie grant agreement, No. 956285 (VAGABOND). This work is part of the research program Vernieuwingsimpuls Vidi (Combining targeted compounds in neuroblastoma tumors; is two better than one?) with project number 91716482, and Veni (Release the beast: Boosting CAR-T cell immunotherapy for neuroblastoma) with project number 09150162010022, which is partly financed by the Dutch Research Council (NWO). In addition, this project has received funding from the European Research Council (ERC) under the European Union’s Horizon 2020 research and innovation program under grant agreement No 716079 Predict, from grant H2020-iPC-826121, and from Villa Joep. The three in vivo studies studying anti-TIGIT/PD-L1 were funded by Hoffman-La Roche. In vivo work at ICH was funded by Hoffman-La Roche and by Stand up to Cancer/CRUK grant RT6188. JA is further supported by supported by GOSH NIHR BRC. The single cell genomics facility of the Princess Máxima Center is funded by KiKa.

