## Supplemental files for "Integrative analysis of neuroblastoma by single-cell RNA sequencing identifies the NECTIN2-TIGIT axis as a target for immunotherapy"

### SUPPLEMENTARY DATA

| | | DEMOGRAPHICS | | | TUMOR STAGE | | | | | SURVIVAL | | TUMOR SAMPLE | |
|---|---|---|---|---|---|---|---|---|---|---|---|---|---|
| **ID** | **Sample ID** | **Sex** | **Age at Dx** | **Age at sampling** | **Risk** | **INRGSS** | **INSS** | **CDK4 amp** | **MYCN amp** | **Survival** | **Survival (months)** | **Tumor (sample) location** | **Months Dx-sampling** |
| **Paired samples** | | | | | | | | | | | | | |
| 1 | 000CGH | F | 4.7 | 4.7 | HR | M | 4 | Non-amp | Non-amp | Alive | 32.57 | left adrenal gland | 0 |
| 1 | 000HKP | F | 4.7 | 5.7 | HR | M | 4 | Non-amp | Non-amp | Alive | 32.57 | left adrenal gland | 12.8 |
| 2 | 000GGU | F | 1.1 | 1.1 | HR | M | 4 | Non-amp | Amp | Deceased | 13.87 | Lymph node left inguinal | -0.23 |
| 2 | 000AKW | F | 1.1 | 1.5 | HR | M | 4 | Non-amp | Amp | Deceased | 13.87 | left adrenal gland | 4.4 |
| 3 | 000BNB | F | 4.0 | 4.0 | HR | M | 4 | Non-amp | Non-amp | Alive | 16.37 | left adrenal gland | 0.03 |
| 3 | 000GXF | F | 4.0 | 4.4 | HR | M | 4 | Non-amp | Non-amp | Alive | 16.37 | left adrenal gland | 4.47 |
| 4 | 000FQM | F | 4.3 | 4.3 | HR | M | 4 | Non-amp | Non-amp | Alive | 22.87 | left cervical primary tumor | -0.03 |
| 4 | 000GWJ | F | 4.3 | 4.6 | HR | M | 4 | Non-amp | Non-amp | Alive | 22.87 | left cervical primary tumor | 3.03 |
| 5 | 000GZZ | F | 12.6 | 12.6 | HR | M | 4 | Non-amp | Non-amp | Alive | 21.23 | right adrenal gland | 0.03 |
| 5 | 000CTN | F | 12.6 | 13.0 | HR | M | 4 | Non-amp | Non-amp | Alive | 21.23 | right adrenal gland | 4.63 |
| **Unpaired pre-treatment samples** | | | | | | | | | | | | | |
| 6 | NB125 | M | 1.2 | 1.1 | OG | L2 | 3 | Non-amp | Non-amp | Alive | 39.33 | left adrenal gland | -0.37 |
| 7 | 000HLR | F | 5.1 | 5.9 | HR | M | 4 | Amp | Non-amp | Alive | 29.87 | left adrenal gland | 10.17 |
| 8 | 000HWF | F | 4.3 | 4.3 | HR | M | 4 | Non-amp | Amp | Alive | 18.83 | right cervical primary | 0.03 |
| 9 | 000AMC | F | 0.8 | 0.8 | HR | M | 4 | Non-amp | Non-amp | Deceased | 12.77 | left adrenal gland | 0.03 |
| 10 | 000HIZ | M | 11.3 | 11.3 | HR | M | 4 | Non-amp | Amp | Alive | 19.7 | right adrenal gland | -0.07 |
| **Unpaired post-treatment samples** | | | | | | | | | | | | | |
| 14 | NB060 | M | 4.0 | 4.4 | HR | M | 4 | Non-amp | Amp | Deceased | 26.13 | Right adrenal gland | 4.5 |
| 17 | NB098 | M | 3.9 | 4.8 | HR | M | 4 | Non-amp | Non-amp | Alive | 50.43 | Left adrenal gland | 10.8 |
| 11 | NB106 | M | 1.3 | 2.7 | MR | L1 | 3 | Non-amp | Non-amp | Alive | 56.9 | left adrenal gland | 16.13 |
| 12 | NB107 | M | 2.3 | 2.7 | HR | M | 4 | Non-amp | Non-amp | Alive | 46.83 | paravertebral lesion left | 4.93 |
| 15 | NB124 | M | 2.5 | 2.8 | HR | M | 4 | Non-amp | Amp | Alive | 43.23 | left adrenal gland | 3.3 |
| 13 | NB130 | M | 4.6 | 5.0 | HR | M | 4 | Non-amp | Non-amp | Deceased | 35.27 | left adrenal gland | 4.63 |
| 19 | NB132 | M | 4.7 | 5.1 | HR | M | 4 | Non-amp | Amp | Deceased | 27.63 | left adrenal gland | 4.87 |
| 16 | NB138 | M | 0.3 | 15.5 | NA | Ganglioneuroma | NA | Non-amp | Non-amp | Alive | 213.37 | inguinal lymph node | 182.07 |
| 18 | NB151 | M | 2.1 | 3.1 | HR | M | 4 | Non-amp | Non-amp | Deceased | 19.23 | left adrenal gland | 11.73 |
| 20 | NB152 | M | 2.0 | 2.4 | HR | M | 4 | Non-amp | Non-amp | Deceased | 27.83 | left adrenal gland | 4.7 |
| **Overall** | | | | | | | | | | | | | |
| | *Median (IQR)* | 8 F | *3.96 (1.83-4.6)* | *4.4 (2.68-5.07)* | 17 HR | 17 M | 17 st. 4 | 1/20 Amp | 6/20 Amp | 7 deceased | *27.7 (19.6-40.3)* | 19/25 adrenal gland | Pre: *0.02 (-0.06-0.03)* Post: *4.7 (4.5-11.3)* |

**Supplementary Table 1. Clinical characteristics of patients.** Tx = treatment, Dx = Diagnosis, INRG = International Neuroblastoma Risk Group, INRGSS = International Neuroblastoma Risk Group Staging System, INSS = International Neuroblastoma Staging System, N5 = cisplatin, etoposide, and vindesine; N6=vincristine, dacarbazine, ifosfamide, and doxorubicine; N8 = topotecan, cyclophosphamide, and etoposide; BuMel = busulfan and melphalan hydrochloride; TEMIRI = temozolomide and irinotecan; MIBG Tx = 131I-Metaiodbenzylguanidine; TOTEM = topotecan and temozolomide.

| ID | Sample ID | Treatment | Treatment until sampling | Months last chemo - sampling | Response to immunotherapy |
|---|---|---|---|---|---|
| | | CHEMOTHERAPY | | | IMMUNOTHERAPY |
| **Paired samples** | | | | | |
| 1 | 000CGH | Pre | untreated diagnostic sample | NA | No event during/after IT |
| 1 | 000HKP | Post | 3x N5N6, 3x TEMIRI, 2x MIBG Tx + topotecan, BuMel | 2 | No event during/after IT |
| 2 | 000GGU | Pre | untreated diagnostic sample | NA | Relapse after 3rd course of IT |
| 2 | 000AKW | Post | 3x N5/N6 | 1 | Relapse after 3rd course of IT |
| 3 | 000BNB | Pre | untreated diagnostic sample | NA | No event during/after IT |
| 3 | 000GXF | Post | 3x N5/N6 | 1 | No event during/after IT |
| 4 | 000FQM | Pre | untreated diagnostic sample | NA | No event during/after IT |
| 4 | 000GWJ | Post | 2x N5/ N6 | 1 | No event during/after IT |
| 5 | 000GZZ | Pre | untreated diagnostic sample | NA | Progression after induction therapy, no event during/after IT |
| 5 | 000CTN | Post | 3x N5N6, 6x TOTEM, 2x MIBG Tx + topotecan | 1 | Progression after induction therapy, no event during/after IT |
| **Unpaired pre-treatment samples** | | | | | |
| 6 | NB125 | Pre | untreated diagnostic sample | NA | No IT (medium risk) |
| 7 | 000HLR | Pre | untreated diagnostic sample | NA | IT after early relapse (last two courses w/o anti-GD2 due to side effects) |
| 8 | 000HWF | Pre | untreated diagnostic sample | NA | No event during/after IT |
| 9 | 000AMC | Pre | untreated diagnostic sample | NA | No IT (deceased before start IT) |
| 10 | 000HIZ | Pre | untreated diagnostic sample | NA | No event during/after IT |
| **Unpaired post-treatment samples** | | | | | |
| 14 | NB060 | Post | 3x N5, 1x N6 | 1 | Relapse after 6th course IT |
| 17 | NB098 | Post | 2x N5N6, 4xN8, BuMel | 1 | No event during/after IT |
| 11 | NB106 | Post | 2xN5, 1xN6 | 1 | No IT (medium risk) |
| 12 | NB107 | Post | 3x N5N6 | 1 | CR after BEACON (anti-GD2+TOTEM) due to relapse before start of regular IT |
| 15 | NB124 | Post | 3x N5N6 | 1 | IT wo IL-2 (liver toxicity) |
| 13 | NB130 | Post | 3x N5N6 | 1 | Therapy related MDS 18 months after IT |
| 19 | NB132 | Post | 3x N5N6 | 1 | Relapse shortly after IT |
| 16 | NB138 | Post | COG MRG protocol, 2x MIBG, isotretinoin. Last Tx 2005 | 156 | No IT (GN) |
| 18 | NB151 | Post | 3x N5N6, 3x TEMIRI, 2x MIBG Tx + topotecan, BuMel | 2 | Progression during IT (during 2nd round) |
| 20 | NB152 | Post | 3x N5N6 | 1 | Progression during IT (during 3rd round) |
| **Overall** | | | | | |
| | *Median (IQR)* | 10/25 Pre | 10/25 untreated, 15/25 chemotherapy | *1 (1-1)* | |

**Supplementary Table 1 (continued). Clinical characteristics of patients.** Tx = treatment, Dx = Diagnosis, INRG = International Neuroblastoma Risk Group INRGSS = International Neuroblastoma Risk Group Staging System, INSS = International Neuroblastoma Staging System, N5 = cisplatin, etoposide, and vindesine; N6=vincristine, dacarbazine, ifosfamide, and doxorubicine; N8 = topotecan, cyclophosphamide, and etoposide; BuMel = busulfan and melphalan hydrochloride; TEMIRI = temozolomide and irinotecan; MIBG Tx = 131I-Metaiodbenzylguanidine; TOTEM = topotecan and temozolomide.

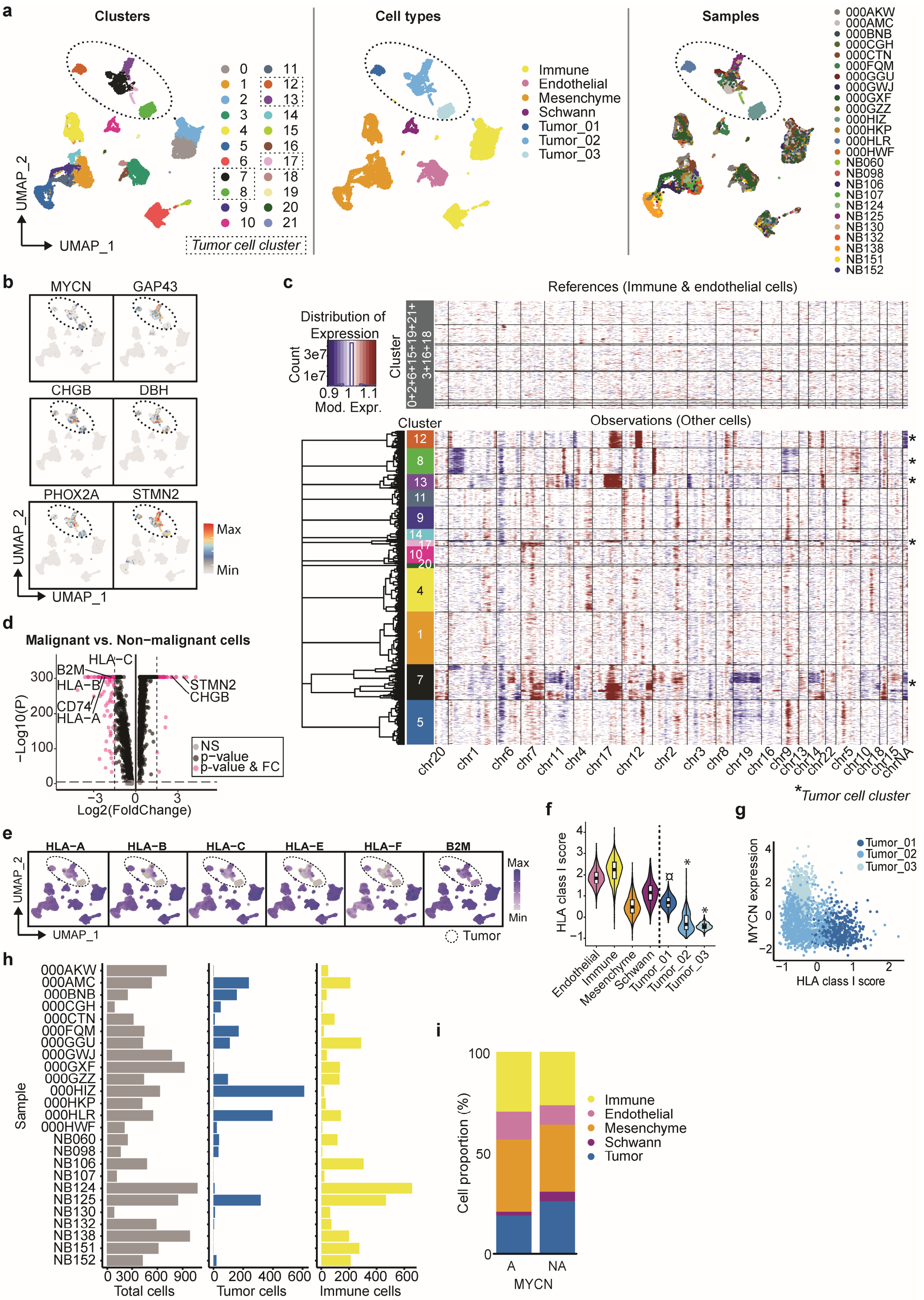


**Supplementary Fig. 1. The single-cell landscape of neuroblastoma. (a)** UMAP of identified cell clusters, main cell types and sample distribution. Dotted circles highlight the tumor cell clusters. **(b)** UMAP with expression of typical neuroblastoma genes (extension of Fig. 1d). **(c)** Copy number profile of the samples, generated using *inferCNV*, to confirm tumor cell identity. The identified cell clusters (panel a) were used as cell groups, in which immune and endothelial cell clusters were used as reference. *Clusters annotated as tumor cells based on gene expression profile. **(d)** Volcanoplot of differentially expressed genes (padj<0.05) between tumor cells (malignant) and all other cells (non-malignant). **(e)** Featureplots of HLA class I genes used to construct the HLA class I gene score. **(f)** Violin plot showing the modulescore of HLA class I genes *HLA-A*, *HLA-B*, *HLA-C*, *HLA-E*, *HLA-F* and *B2M* in the main cell types. *Kruskall-Wallis + Dunn's. °p<0.0001 versus all healthy clusters except mesenchyme (ns). *p<0.0001 versus all healthy clusters.* **(g)** Correlation between *MYCN* expression and HLA class I gene score for the three tumor clusters. **(h)** Number of total cells, tumor cells and immune cells per sample. **(i)** Cell proportion in MYCN-A versus MYCN-NA samples.

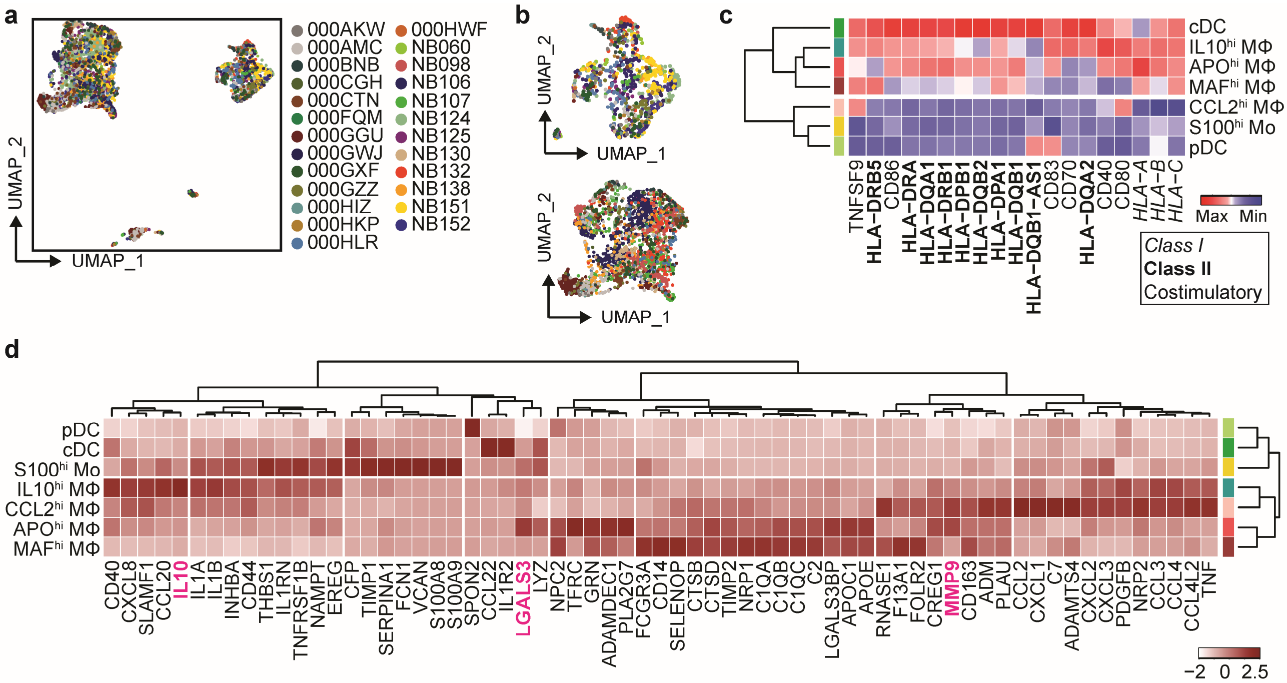


**Supplementary Fig. 2. The immune environment of neuroblastoma. (a)** UMAP of sample distribution in immune compartment. **(b)** UMAP of sample distribution in myeloid cell (*upper*) and lymphoid cell (*lower*) compartments. **(c)** Heatmap of genes representing antigen presenting/co-stimulatory capacity. Genes are included in module score shown in Fig. 2c. **(d)** Heatmap with secreted factors which were 1) specifically upregulated in myeloid cells compared to all other cells and 2) among the differentially expressed genes between the myeloid clusters.

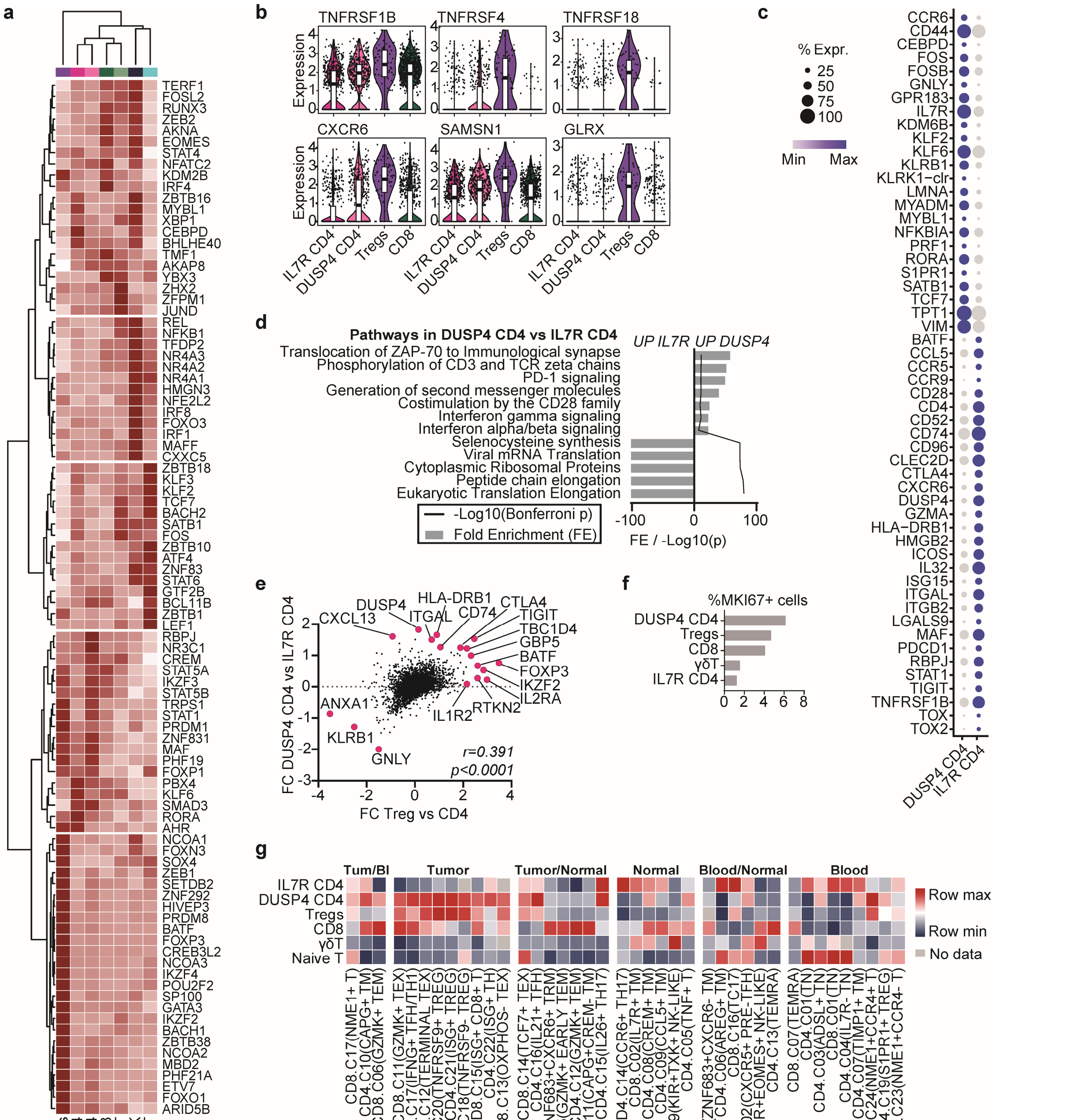

**Supplementary Fig. 3. Neuroblastoma is characterized by immunosuppressive and dysfunctional lymphoid populations. (a)** Heatmap of transcription factors that were among the differentially expressed (padj<0.05) genes between T/NK cell subclusters. **(b)** Expression of effector Treg marker genes which are significantly upregulated (padj<0.05) in neuroblastoma-infiltrating Tregs. **(c)** Dotplot with selected differentially expressed genes (padj<0.05) between the two CD4 T effector populations. **(d)** Pathway analysis (Reactome; Bonferroni-corrected p-value) of differentially expressed genes (padj<0.05) between the *DUSP4*$^{hi}$ and *IL7R*$^{hi}$ CD4 T cell populations. **(e)** Correlation between differentially expressed genes in *DUSP4*$^{hi}$ versus *IL7R*$^{hi}$ CD4 T cells, and Tregs versus *DUSP4*$^{hi}$ CD4 T cells. *FC = fold change.* **(f)** Percentage of T cells expressing proliferation marker *MKI67*. **(g)** GSEA of neuroblastoma immune cell clusters (horizontal) with previously published gene signatures(40) which identify T cell clusters specifically associated with blood, normal tissue and tumors (vertical), showing correspondence between the cluster signatures.

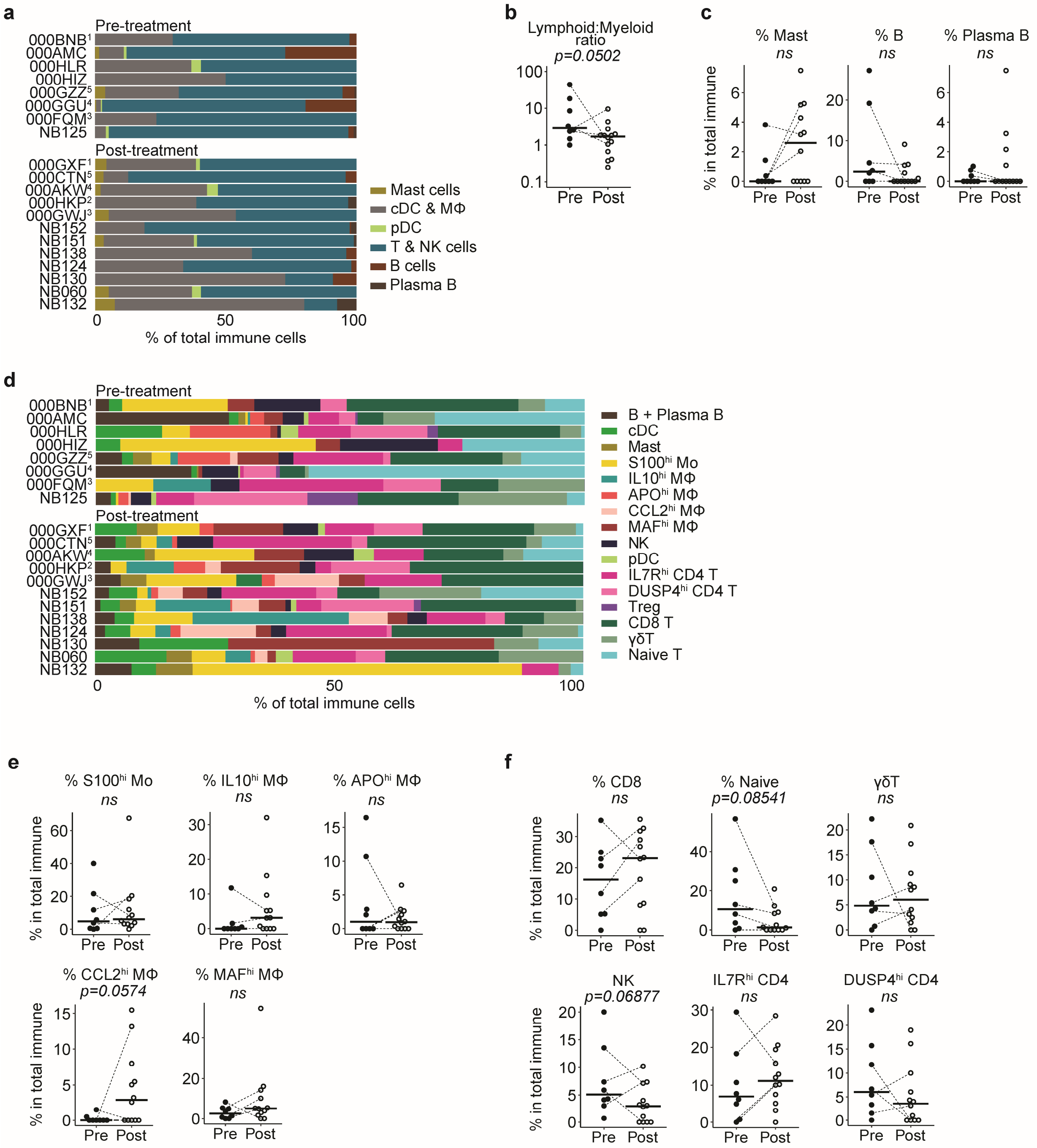

**Supplementary Fig. 4. The immune cell composition in neuroblastoma before and after induction chemotherapy. (a)** Composition of overall immune cells per patient before and after induction chemotherapy. [1,2,3,4,5]: Paired samples before and after treatment. **(b)** Lymphoid:myeloid ratio before and after chemotherapy. Dotted lines indicate paired samples. *Mann-Whitney U test.* **(c)** Mast cell, B cell and plasma B cell percentages before and after induction chemotherapy. Dotted lines indicate paired samples. *Mann-Whitney U test.* **(d)** Composition of detailed immune cell clusters per patient before and after induction chemotherapy. [1,2,3,4,5]: Paired samples before and after treatment. **(e)** Myeloid cell percentages before and after induction chemotherapy. Dotted lines indicate paired samples. *Mann-Whitney U test.* **(f)** Lymphoid cell percentages before and after induction chemotherapy. Dotted lines indicate paired samples. *Mann-Whitney U test.*

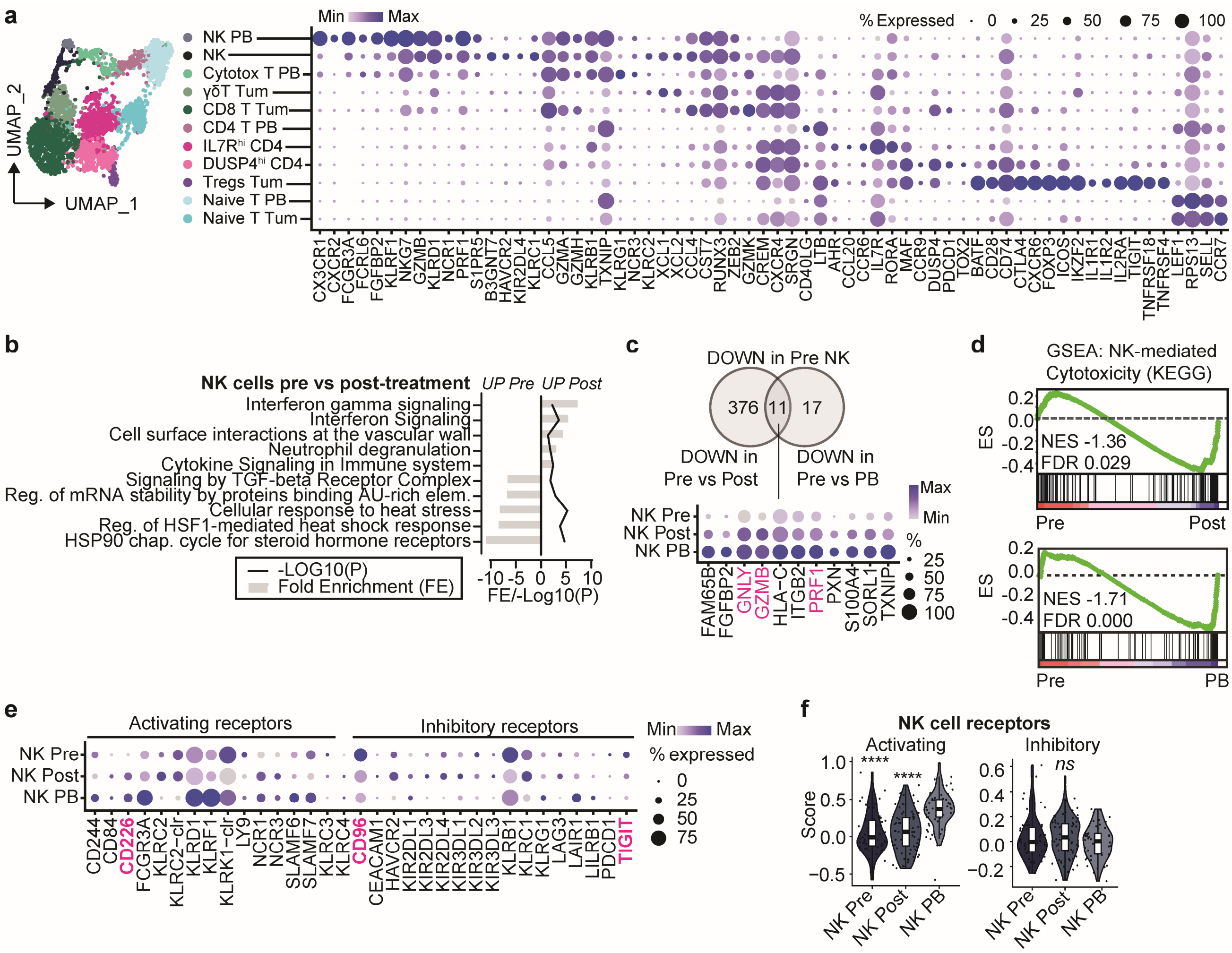

**Supplementary Fig. 5. NK cells in neuroblastoma show reduced cytotoxicity pre-treatment. (a)** UMAP and dotplot of T/NK subsets from both tumor and reference peripheral blood (PB) of adult healthy donors showing a selection of their differentially expressed genes. **(b)** Pathway analysis of NK cells pre- versus post-treatment (Reactome, bonferroni-corrected p-values<0.05). **(c)** Significantly downregulated genes in NK cells in pre-treatment tumors compared to either NK cells post-treatment (nominal p<0.05) or reference peripheral blood (PB) NK cells (padj<0.05). **(d)** Gene set enrichment analysis (GSEA) of NK cells in pre-treatment tumors compared to NK cells from post-treatment tumors and peripheral blood (PB). *NES=normalized enrichment score, FDR=false discovery rate.* **(e)** Dotplot of NK cell activating and inhibitory receptor expression. **(f)** Modulescore of NK cell activating and inhibitory receptors shown in Supplementary Fig. 5e. *Kruskall-Wallis with Dunn's, ****P<0.0001 vs NK PB, ns=not significant*.

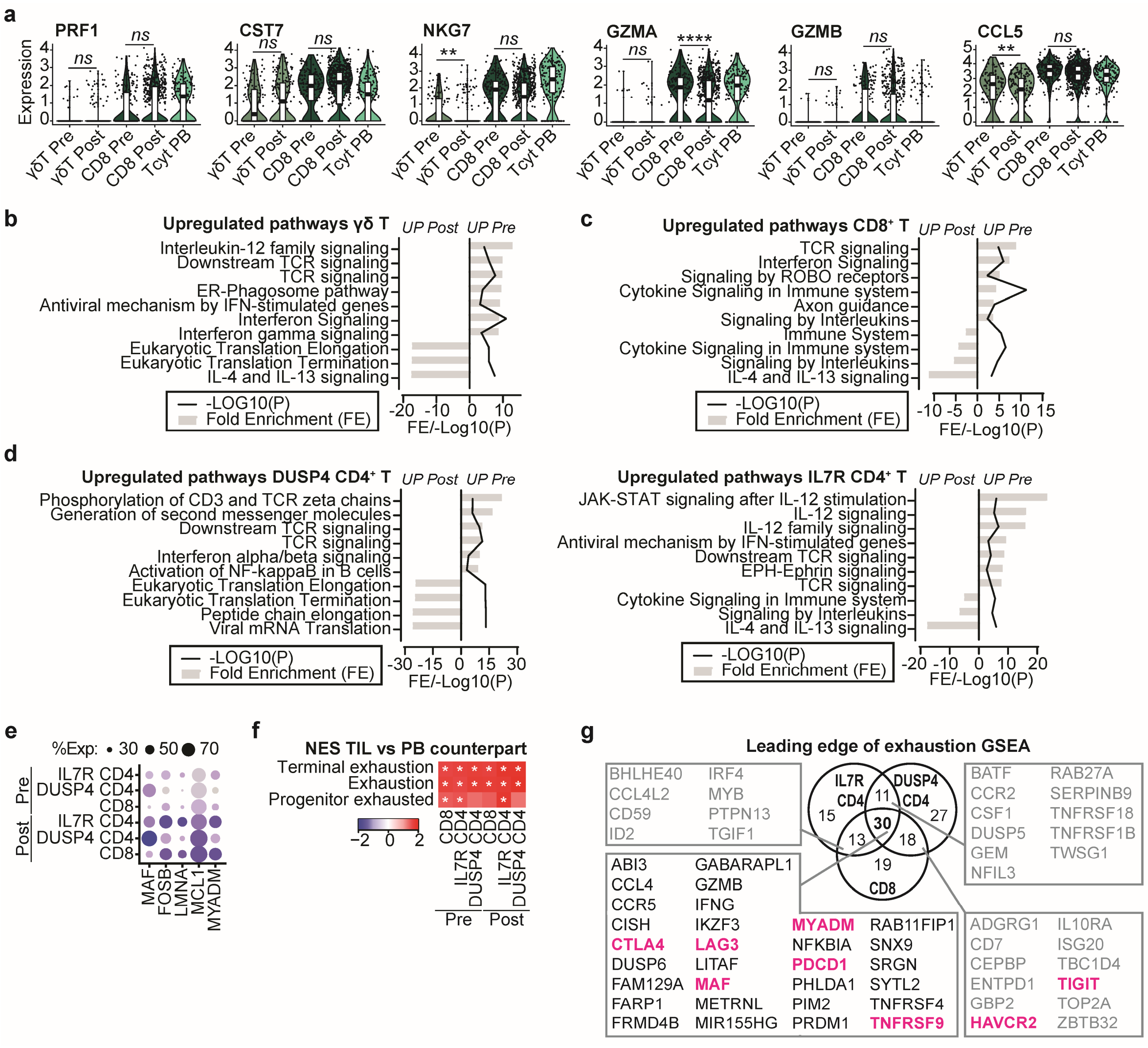

**Supplementary Fig. 6. T cells in neuroblastoma show increased dysfunctionality post-treatment. (a)** Violin plots of genes associated with cytotoxicity in γδ T and CD8 T cells pre- and post-treatment compared to cytotoxic T cells in reference blood (Tcyt PB). ***Nominal p<0.05 pre vs post. ****Nominal p<0.0001 pre vs post*. **(b-d)** Pathway analysis of differentially expressed pathways in γδ T **(b)**, CD8 T **(c)** and CD4 T **(d)** pre- versus post-treatment (Reactome, bonferroni-corrected p-values<0.05). **(e)** Dotplot of shared upregulated genes comparing expression in CD4 and CD8 T cells pre- and post-treatment. **(f)** Gene set enrichment analysis (GSEA) of exhaustion signatures (terminal & progenitor: GSE84105, exhaustion: Man et al.(49)) in indicated CD4 and CD8 T cell clusters versus their respective counterparts from reference blood. **(g)** Venn diagram of shared upregulated genes post-treatment versus pre-treatment, which are present in leading edges (core enriched) of GSEA analysis with exhaustion/dysfunction signatures from Fig. 4m.

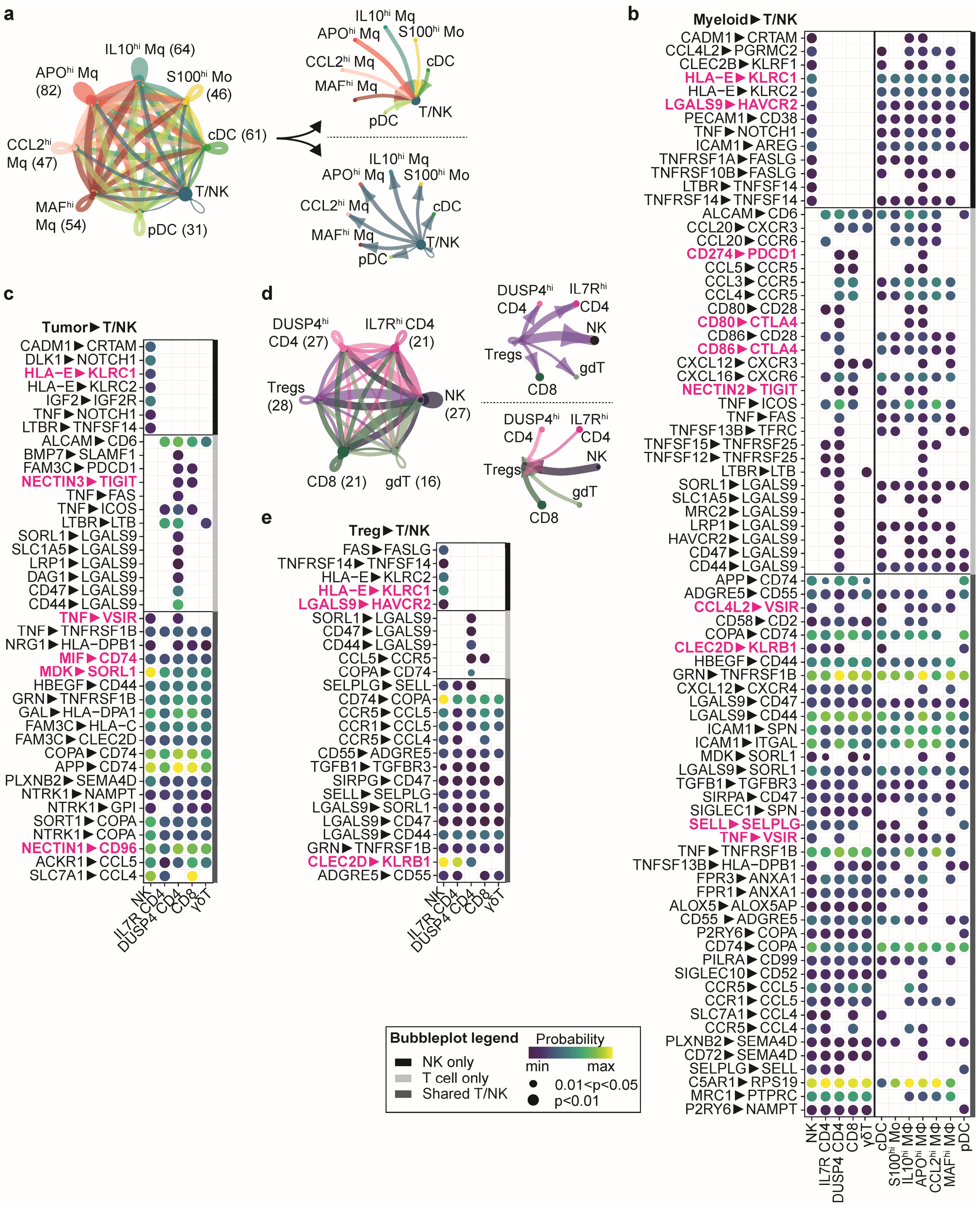


**Supplementary Fig. 7. Immunoregulatory interactions in the neuroblastoma tumor-microenvironment. (a)** Interaction network involving interactions between myeloid cells and T/NK cells constructed with CellChat(50). **(b)** Bubbleplot of predicted interactions between myeloid cells and indicated T/NK subsets. Interactions with each specific T/NK subset and each specific myeloid subset were evaluated and subsequently merged, with the highest probability of each interaction pair depicted in the plot (*left: myeloid merged; right: T/NK merged*) **(c)** Bubbleplot of predicted interactions between tumor cells and indicated T/NK subsets. **(d)** Interaction network of interactions among Tregs and all other T/NK cell subsets. **(e)** Bubbleplot involving interactions between Tregs and indicated T/NK cell subsets.

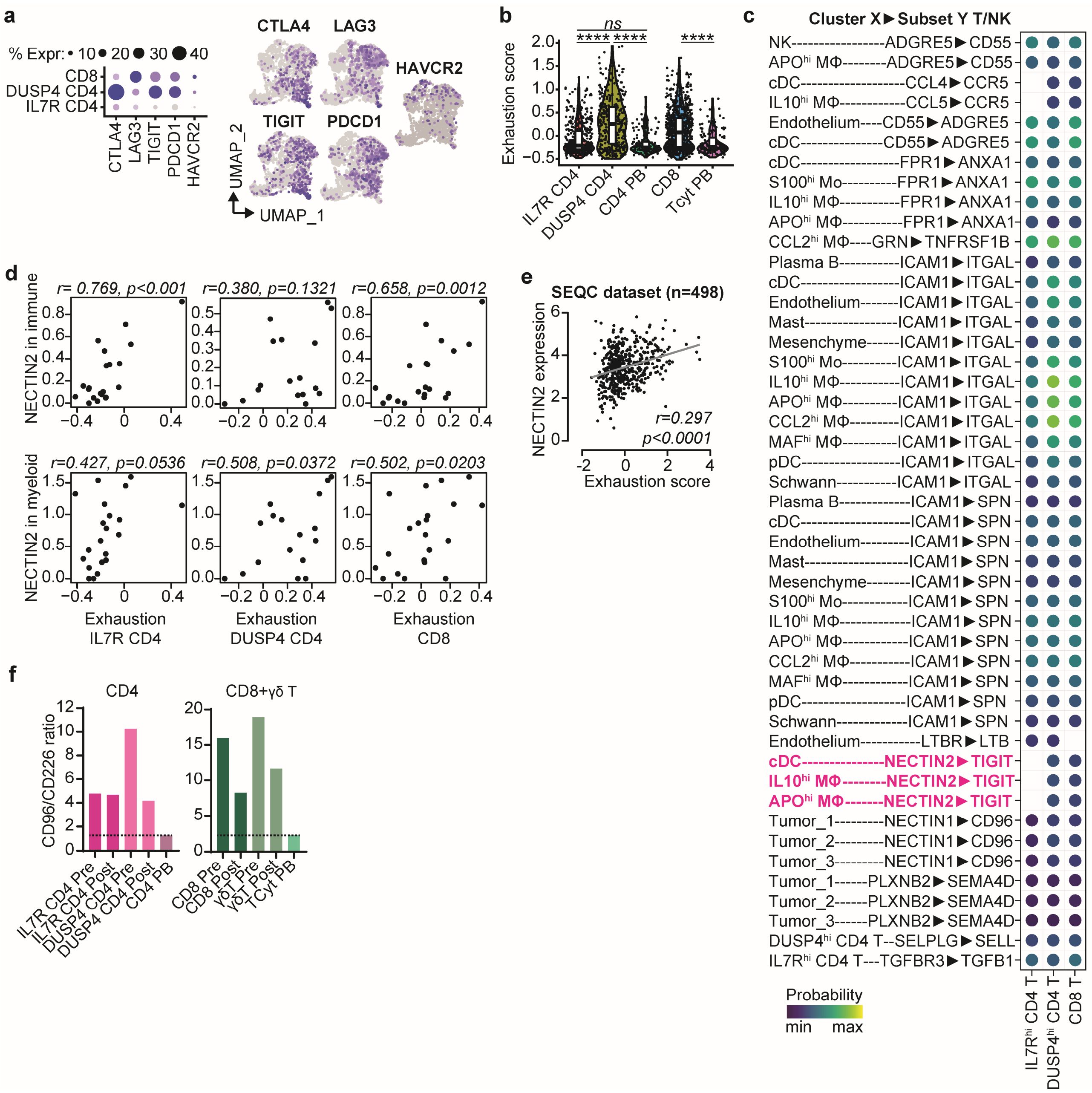

**Supplementary Fig. 8. The *NECTIN2-TIGIT* axis is associated with T cell dysfunction. (a)** Dotplot with expression of dysfunction/exhaustion markers *CTLA4*, *LAG3*, *TIGIT*, *PDCD1* and *HAVCR2* in the indicated T cell subsets and Featureplots showing the log-normalized expression of these genes in the T/NK cell compartment. **(b)** Modulescore of the dysfunction/exhaustion genes in the indicated T cell subsets. *Kruskall Wallis + Dunn's.* **(c)** Bubbleplot showing detailed interaction analysis involving the 13 genes that showed the highest positive correlation with either *DUSP4*$^{hi}$ CD4, *IL7R*$^{hi}$ CD4 or CD8 T cell dysfunction/exhaustion. **(d)** Correlation of *NECTIN2* expression in immune and myeloid cells with exhaustion scores of *DUSP4*$^{hi}$ CD4, *IL7R*$^{hi}$ CD4 and CD8 T cells. **(e)** Correlation of *NECTIN2* with exhaustion score in bulk-RNAseq dataset of SEQC cohort consisting of 498 neuroblastomas (r2.amc.nl; Tumor Neuroblastoma - SEQC - 498 - RPM - seqcnb1; GSE49710). **(f)** *CD96*/*CD226* ratio in T cell subsets pre-/post-treatment and in reference blood (PB).

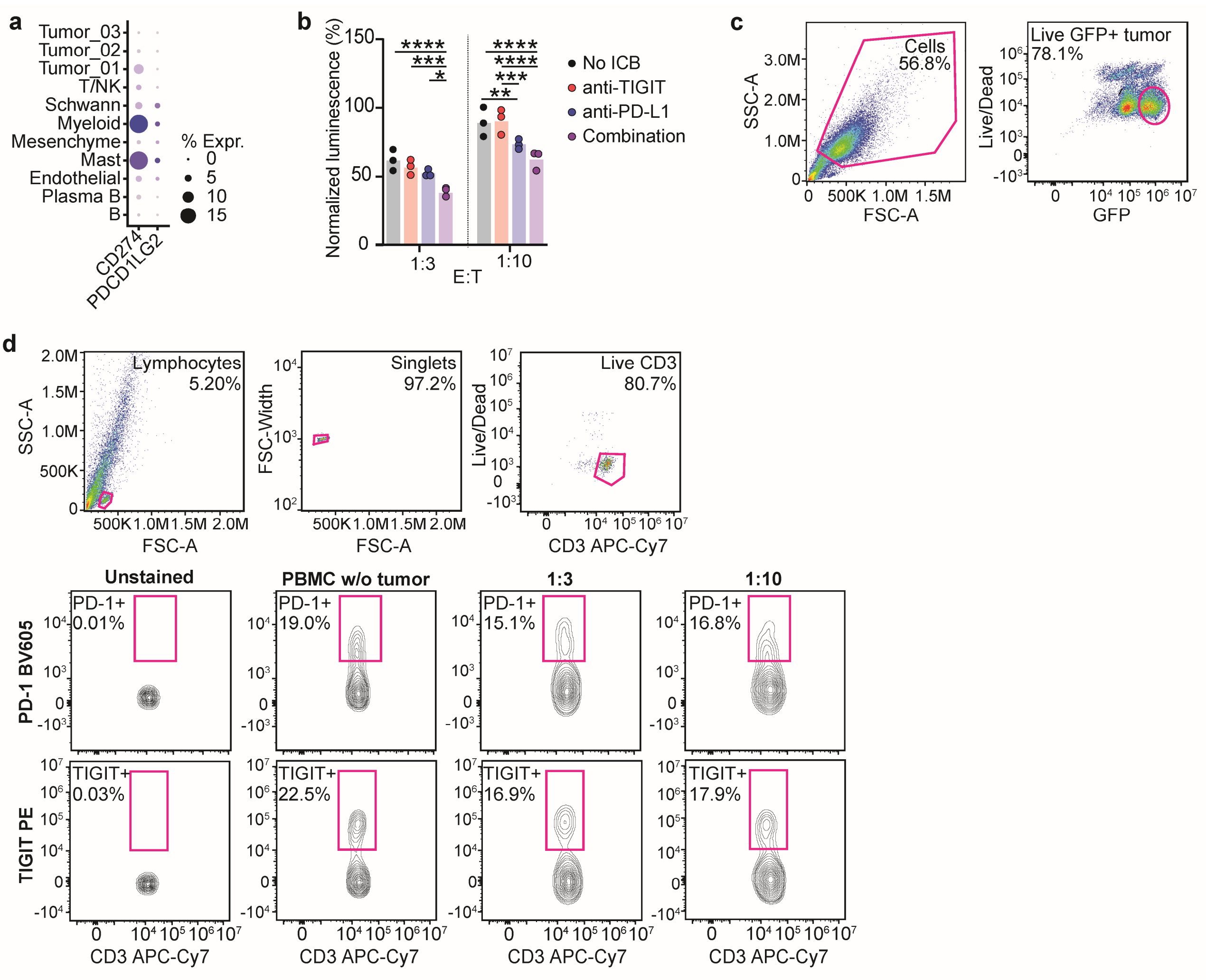


**Supplementary Fig. 9. Combined TIGIT/PD-L1 blockade enhances immune responses against neuroblastoma *in vitro*. (a)** Dotplot with expression of PD-1 ligands *CD274* (PD-L1) and *PDCD1LG2* (PD-L2) in neuroblastoma. **(b)** Luminescence readout of *in vitro* tumor killing assay with neuroblastoma (luciferase-transduced) tumoroids and healthy donor PBMC at effector:target ratios 1:3 and 1:10. Tumor cells and PBMC were cocultured for 6 days with or without anti-TIGIT and/or anti-PD-L1. Luminescence values were normalized against condition with only untreated tumoroids. *n=3. Two-way ANOVA with Tukey. #p<0.0001 vs No PBMCS-No ICB; *p<0.05; **p<0.01, p<0.001, p<0.0001*. **(c)** Gating strategy for flow cytometric expression analysis of Nectin-2, PVR and PD-L1 expression GFP-positive tumoroids as shown in Fig. 6c. **(d)** Flow cytometric analysis of PD-1 and TIGIT expression on PBMC cocultured with neuroblastoma tumoroids for 6 days. PBMC w/o tumor was measured at timepoint 0 before coculture.

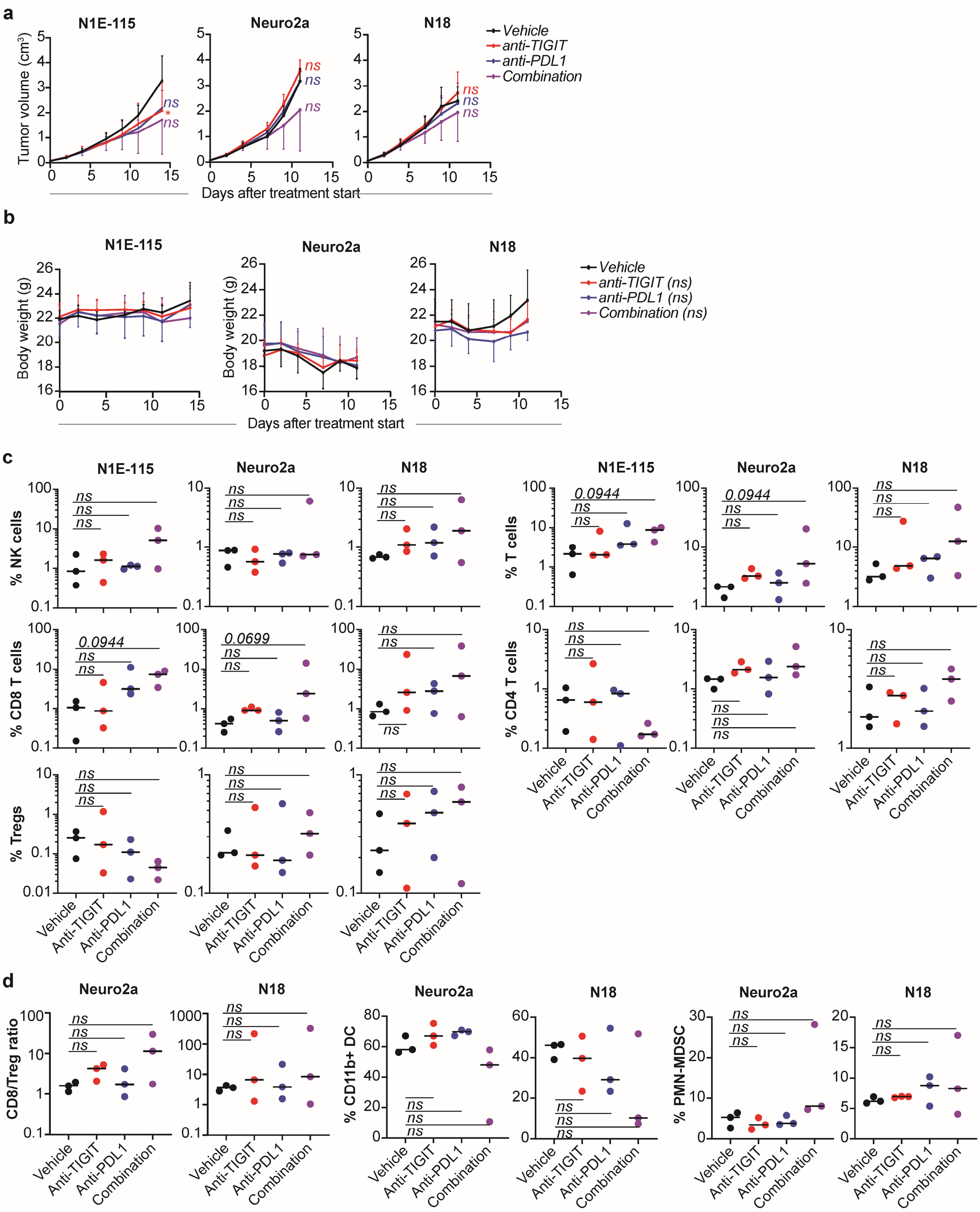


**Supplementary Fig. 10. Combined TIGIT/PD-L1 blockade enhances immune responses against neuroblastoma *in vivo*. (a)** Tumor volume in N1E-115, Neuro2a and N18 mouse models (n=6 per group) treated with anti-TIGIT and/or anti-PD-L1 up to day 15 of treatment. **(b)** Body weight of N1E-115, Neuro2a and N18 mouse models (n=6 per group) treated with anti-TIGIT and/or anti-PD-L1 up to day 15 of treatment. **(c)** Flow cytometric analysis of proportion of T/NK cells subsets infiltrating tumors in N1E-115, Neuro2a and N18 models. *Kruskall-Wallis with Dunn's.* **(d)** Flow cytometric analysis of tumor-infiltrating leukocytes in Neuro2a and N18 models (n=3 per group) showing CD8/Treg ratio (*left*) fraction of and CD11b+ dendritic cells (*middle*) and polymorphonuclear myeloid-derived suppressor cells (PMN-MDSC) (*right*) in mice treated with anti-TIGIT and/or anti-PD-L1 for 7 days. *Kruskall-Wallis with Dunn's.*

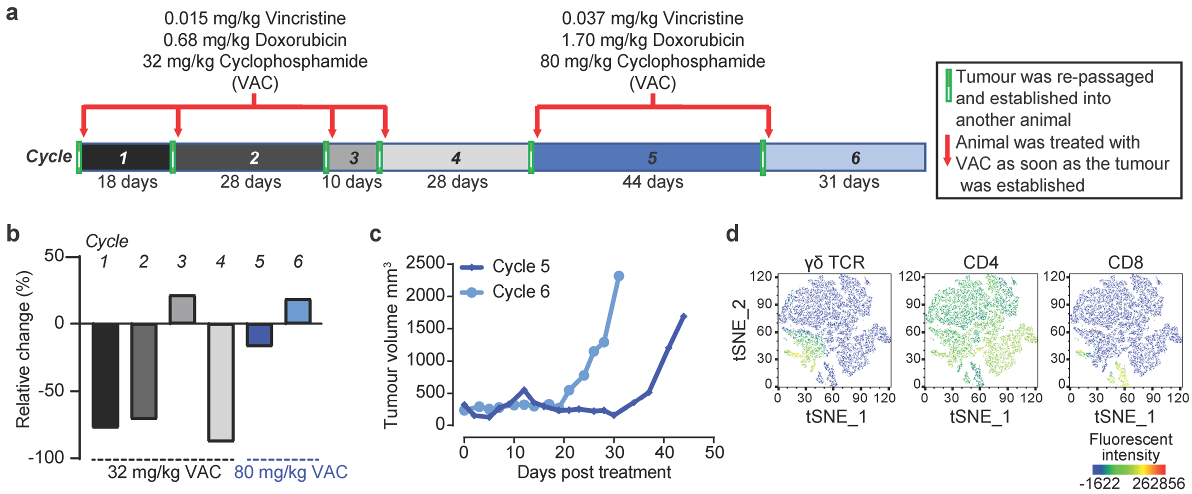


**Supplementary Fig. 11. Generation of the VAC (Vincristine, Adriamycin/Doxorubicin, Cyclophosphamide) resistant model. (a)** Summary of dose escalation treatment schedule to achieve resistance. Red arrows indicate when single dose VAC was administered. The number of days taken for the tumor to relapse is indicated below each cycle. At the end of each cycle, the allograft tumor was re-passaged into another animal before treatment started. **(b)** Percentage relative tumor volume change 7 days after treatment with single dose of VAC at 32 mg/kg or 80 mg/kg. **(c)** Time taken for allograft tumor to relapse after treatment with single high dose of VAC (80 mg/kg VAC) in Cycle 5 and Cycle 6. Tumor cells after Cycle 6 were used to establish allografts for therapeutic studies. **(d)** Expression of immune cell markers γδ TCR, CD4 and CD8 in $CD45^+$ tumor-infiltrating immune cells by flow cytometry.
